# RNA structure and multiple weak interactions balance the interplay between RNA binding and phase separation of SARS-CoV-2 nucleocapsid

**DOI:** 10.1101/2023.07.02.547440

**Authors:** Aidan B Estelle, Heather M Forsythe, Zhen Yu, Kaitlyn Hughes, Brittany Lasher, Patrick Allen, Patrick Reardon, David A Hendrix, Elisar J Barbar

## Abstract

The nucleocapsid (N) protein of SARS-CoV-2 binds viral RNA, condensing it inside the virion, and phase separating with RNA to form liquid-liquid condensates. There is little consensus on what differentiates sequence-independent N-RNA interactions in the virion or in liquid droplets from those with specific genomic RNA motifs necessary for viral function inside infected cells. To identify the RNA structures and the N domains responsible for specific interactions and phase separation, we use the first 1000nt of viral RNA and short RNA segments designed as models for single-stranded and paired RNA. Binding affinities estimated from fluorescence anisotropy of these RNAs to the two folded domains of N (the NTD and CTD) and comparison to full-length N demonstrate that the NTD binds preferentially to single-stranded RNA, and while it is the primary RNA binding site, it is not essential to phase separation. Nuclear magnetic resonance spectroscopy identifies two RNA binding sites on the NTD: a previously characterized site and an additional although weaker RNA-binding face that becomes prominent when binding to the primary site is weak, such as with dsRNA or a binding-impaired mutant. Phase separation assays of nucleocapsid domains with different RNA structures support a model where multiple weak interactions, such as with the CTD or the NTD’s secondary face promote phase separation, while strong, specific interactions do not. These studies indicate that both strong and multivalent weak N-RNA interactions underlie the multifunctional abilities of N.

**Significance:** The nucleocapsid protein of the SARS-CoV-2 coronavirus binds to viral RNA, both to protect and condense it inside the viral particle and to facilitate viral transcription inside infected host cells. Evidence suggests that variations in RNA structure impact how and where it binds to the nucleocapsid, but these differences are not well understood at a structural level. Using nuclear magnetic resonance spectroscopy, we examine the interactions between each folded domain of the nucleocapsid and different RNA structures. Binding affinities and NMR chemical shift profiles demonstrate that binding between the N-terminal domain and single stranded RNA is driven by strong interactions at a specific site, while multiple weak nonspecific interactions at newly discovered sites lead to phase separation and RNA condensation.

## Introduction

The SARS-CoV-2 coronavirus is a positive sense RNA virus with a genome approximately 30,000 bases in length that encodes 29 proteins, including four primary structural proteins: the envelope, membrane, spike and nucleocapsid^1, 2^. Among viral proteins, the nucleocapsid (N) is the most abundantly expressed within infected cells^3, 4^, and is highly conserved, with over 90% sequence similarity to the nucleocapsid of SARS-CoV. N packages and protects the viral genome in viral particles by condensing genomic RNA (gRNA) inside the mature virion^1, 5^. Besides RNA packaging, the protein also facilitates viral transcription and replication in infected cells through interactions with RNA^6,7,8^, and forms liquid-liquid phase separated droplets with RNA and other viral proteins^9,10,11,12^

Structurally, the nucleocapsid consists of two folded domains linked and flanked by three intrinsically disordered regions (IDRs) (Fig. 1a)^13, 14^. The N-terminal domain (NTD) spanning residues 47 to 174 is a predominantly β structure with flexible loops surrounding a central β sheet^15,16,17^ The β sheet, including a large hairpin, forms a positively charged cup shape that binds to RNA (Fig. 1c)^15, 18, 19^. While both folded domains bind RNA, the NTD is considered the primary RNA-binding site of N, and binds RNA preferentially over the C-terminal domain (CTD)^20^. The CTD spanning residues 255 to 366 is a stable dimer and forms a disc-like shape^17^ with a positively charged α face made of interlocking α helices, and a β face consisting of a single β sheet with swapped strands linking the dimer’s subunits (Fig. 1d). The intrinsically disordered linker between domains binds weakly to RNA^21, 22^, phase separates with it under some conditions^23, 24^, and also increases the NTD’s affinity to RNA^19, 25^. The full length nucleocapsid binds multivalently to the first 1000 nt of viral RNA more tightly than individual domains^13^, and NMR studies clearly show peak attenuation of the NTD residues and not the linkers^13^ further confirming that the NTD is the primary site of RNA binding and suggesting that multivalency is the primary cause for tighter binding of the full length N^13^. Binding to the CTD and the disordered regions could also contribute to the enhanced affinity, and so does the length of the RNA^13, 19, 25^.

**Figure 1:**
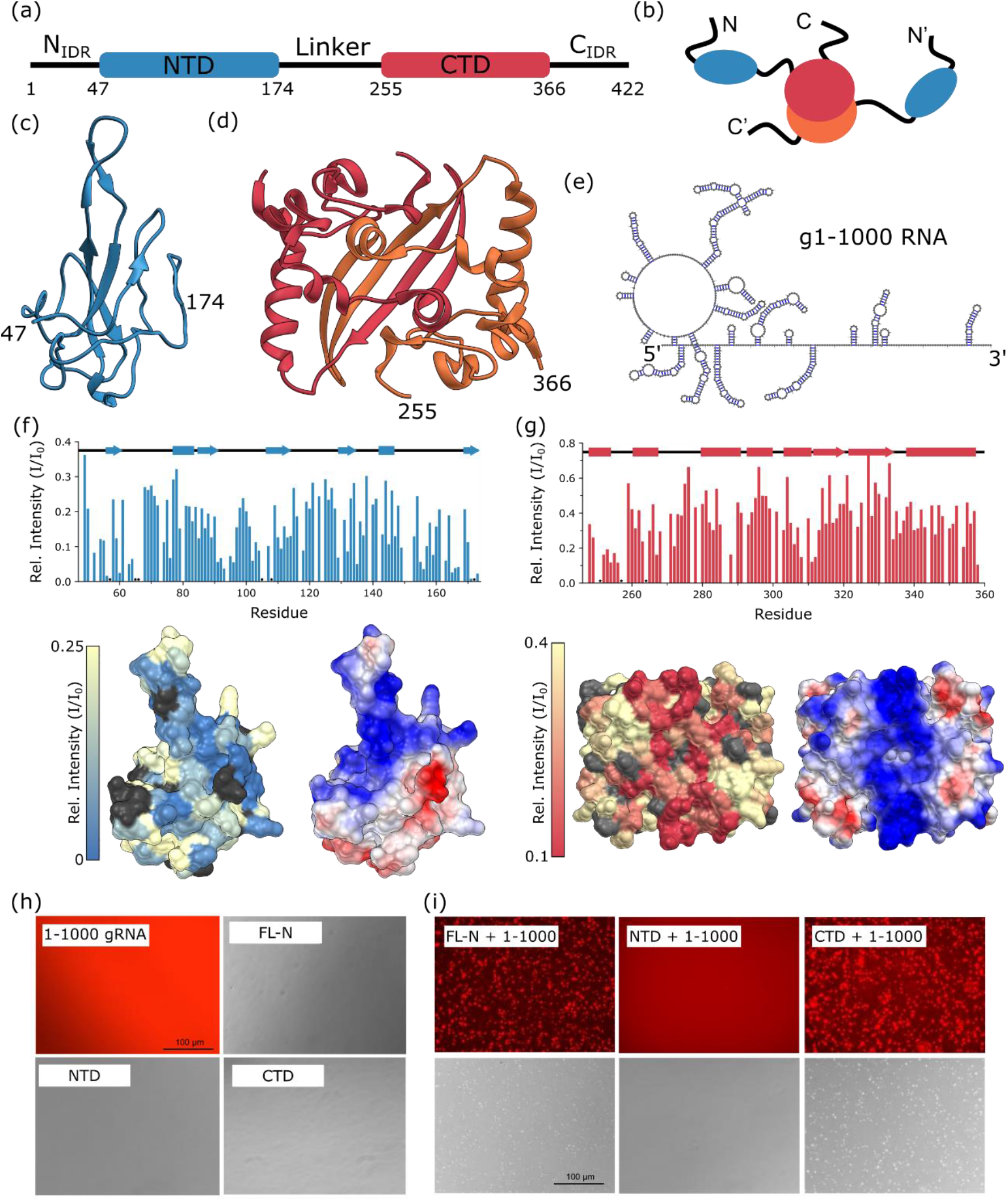
Structure of N and interactions with g1-1000 RNA. (a) Domain diagram of the nucleocapsid protein, showing folded domains (rectangles) and regions of disorder (lines). (b) Cartoon diagram of the dimeric structure of N. (c,d) Ribbon diagrams of NTD (c, PDB 7CDZ) and CTD (d, PDB 7CE0) structures, red and orange denoting different subunits of the dimer structure. (e) Secondary structure for g1-1000, the first 1000 bases of the SARS-CoV-2 nucleocapsid, as determined by SHAPE-MaP measurements^61^. (f,g) Relative peak intensity of the NTD (f) and CTD (g) upon binding to 1-1000 gRNA at a 1:100 RNA:NTD or CTD (top). Relative peak intensities mapped to the NTD surface (bottom left), and surface charge (bottom right). Color scale for surface charge is from -10 (red) to 10 (blue) kcal/(mol*e)^62^. Unassigned and overlapped peaks whose relative peak intensities could not be extracted are colored dark gray. (h) Fluorescence image of free g1-1000 RNA, along brightfield images of each N construct – FL-N, NTD and CTD. (i) Fluorescence (top) and brightfield (bottom) images of FL-N, NTD, and CTD with g1-1000 RNA at 3.6 μM protein and 50 nM g1-1000 at 150 mM NaCl and 37 °C.

N binds RNA both promiscuously and specifically – the protein binds tightly to any RNA regardless of sequence, but it is also recruited to specific motifs within the viral genome. In SARS-CoV – the virus which caused the 2002 SARS outbreak– N recognizes specific sequences within gRNA, such as the highly conserved transcription regulation sequence^26^ and packaging signal^27, 28^ that have been a focus of study in SARS-CoV-2 as well^15, 19, 29, 30^, although their exact role is not yet clear. Crosslinking experiments with viral RNAs^10^ have demonstrated that the nucleocapsid is recruited disproportionately to specific regions of the 5’ UTR of the viral genome. In contrast to this sequence-specific binding, many studies have demonstrated that the nucleocapsid binds to RNA (and DNA^18, 31^) regardless of sequence^10, 15, 29, 32^. Indeed, this binding is important to viral function, as both condensation of viral RNA inside the virion and liquid-liquid phase separation are driven by promiscuous interactions between N and RNA^1, 9^.

While structural details on what differentiates specific binding to sites like those in the 5’ UTR from promiscuous interactions that promote phase separation or encapsidation are of particular interest^15, 18, 19, 29^, the mechanism that drives specific binding and how it differs from promiscuous binding are still unclear. Although the IDRs are involved in RNA binding^21, 22^, specific interactions necessitate complex contacts that depend on tertiary structure, meaning that the N- and CTD are the likely facilitators of specific motif recognition. Investigation is complicated by binding of the multiple domains of the nucleocapsid, and by the variety of viral RNA structures, which include both flexible unpaired ssRNA as well as stem-loops and other regions of paired dsRNA^33^. Cross-linking experiments have indicated that single-stranded (ss) RNA may recruit N preferentially over double-stranded (ds) RNA, and that this preference may drive the pattern of localization of N observed in the 5’ UTR^10^. RNA structure plays a role in phase separation as well, as increasing the double-stranded content of RNA promotes phase separation^34^.

As the protein’s primary RNA-binding domain, the NTD is a likely candidate for driving specific N-RNA binding. There are multiple structural investigations of RNA binding to the NTD^15, 18, 19, 29, 35^. Foundational work by Dinesh et al. (2020)^15^ demonstrated that the NTD binds 7- mers of both single and double stranded RNA at the domain’s charged face, although the authors do not compare the binding affinity between the ss- and dsRNA fragment. A docked model of the interaction indicated that the RNA’s charged backbone faces the NTD, suggesting binding is likely to be of low RNA sequence specificity^15^. Mutations that reversed the NTD’s positive charge weakened binding to RNA, further indicating that charge drives the interaction^15^. Work by Redzic et al. (2021)^19^ examined NTD binding for several RNAs of varied length and structure, from a 7- mer ssRNA to a 30-mer stem loop and showed that the site of binding on the NTD was the same for every RNA, although affinities varied, appearing to correlate with the length and flexibility of the RNA. Notably, the stem loop bound the NTD less strongly than a 12-mer ssRNA, although the mix of secondary structure in a stem loop makes it difficult to determine whether the NTD favors binding to ss- or dsRNA^19^. We and others have proposed a role for entropy and dynamics in RNA selection^29, 36^. Very recently, Korn et al^29^ demonstrated that some viral RNA sequences appear to bind the NTD in a distinct mode that is mediated through both electrostatics and conformational stability. In addition, NTD favors RNAs with labile SLs and disfavors pure dsRNA and stable SL elements, suggesting that overall, the NTD has a weak preference for regions of unpaired RNA^29^.

The CTD is also known to bind RNA^37,38,39^, however much less information on CTD-RNA interaction is available, as the interaction was overlooked in early investigations of the nucleocapsid. Gel shift and fluorescence anisotropy assays have demonstrated that the CTD binds to short oligomers of ssRNA, ssDNA and dsDNA^25, 38^, as well as long fragments of viral genomic RNA^13^. A recent study measured binding between the CTD and short 7-mers of RNA and DNA by NMR, finding a cluster of residues towards the N-terminus of the domain with large chemical shift perturbations, although in general the interaction was very weak^20^. A study examining a stem-loop from the 3’ UTR containing both paired and unpaired bases recapitulated this result, and additionally generated a docked model of the RNA bound to the CTD^40^. The model indicated that the RNA sits at the edge of the CTD, binding to a helix at the periphery of the disk-shaped CTD and additionally making contact at several places on the β sheet^40^. Interestingly, the model indicated that paired stems are the primary point of contact on the RNA^40^. While these studies provide a wealth of new information on how the CTD binds RNA, no direct comparison between ssRNA and dsRNA has been carried out, and thus it is not clear how RNA structure impacts affinity or specificity in binding between the CTD and RNA.

With a focus on the NTD and CTD we examine binding to the first 1000 nucleotides of the CoV-2 genomic RNA (g1-1000) and to short 14-nucleotide RNAs designed to form pure single- and double-stranded RNA species. We find that the NTD binds tightly to ssRNA and weakly to dsRNA, especially in the context of the full length nucleocapsid protein. We demonstrate that an additional face of the NTD interacts weakly with RNA and promotes liquid-liquid phase separation. Further, a mutation designed to abrogate interaction between the NTD and RNA promotes binding at this second face, suggesting that the RNA binding to the NTD is in equilibrium between these two distinct binding modes. With the CTD, we confirm that the protein interacts only weakly with both ss and dsRNA, and that dsRNA binds to the positively charged face of the CTD. Examining the propensity of each domain to form phase separated droplets, we find that weak, nonspecific interactions are the primary drivers of droplet formation, and that while the CTD drives phase separation, binding at the weaker face of the NTD can also promote droplet formation. From these results we propose that both strong and weak interactions with RNA are essential for nucleocapsid function.

## Results

### The nucleocapsid domains bind 1-1000 genomic RNA at their positively charged faces

To map RNA-interacting sites on the nucleocapsid, we previously measured changes in NMR spectra of the nucleocapsid following addition of the first 1000 bases of the coronavirus genomic RNA (g1-1000)^13^. Since spectra of FL-N are missing the CTD peaks preventing analysis of CTD-RNA binding, we perform here similar experiments but on individual domains and demonstrate that the CTD also binds g1-1000. Both RNA-bound spectra exhibit no significant chemical shift perturbation, but experience dips in peak intensities, indicative of an equilibrium with a large bound state. The peak intensity ratios (Fig. 1f,g) report on the proximity of each residue to the site of binding, allowing us to roughly map where binding occurs on the protein.

The NTD binds RNA at the positively charged groove (Fig. 1f) around the domain’s β hairpin. The sharpest decrease in peak intensities corresponds to the base of the β hairpin at residues 91-96 and 101-107, as well as the β strand at the N terminus (Fig.1f). Peak intensities are also low at the C-terminus of the domain (residues 160-170), which is not positively charged, but is next to the charged patch in the structure. This pattern of interaction agrees well with published NMR studies of RNA-protein interactions of the NTD^15, 19^, and docked models of the NTD-RNA complex which both indicate that the RNA-NTD complex forms at the charged surface of the NTD^15^, and that RNA wraps around the surface making contacts with the C-terminal end of the domain.

The CTD binds g1-1000 RNA along its positively charged α face (Fig. 1g). These spectra were collected in high (>200 mM) salt concentrations to improve solubility reduce formation of liquid-liquid phase separated droplets with RNA. In the spectrum of the bound CTD, peak intensities are lowest at the N-terminus of the CTD (residues 247-270), as well as at the helical region 305-314 (Fig. 1g). Both the N-terminal region and the helical region around residue 310 are on the α-helical face of the domain, indicating that binding occurs along this face.

### The NTD alone does not phase separate with g1-1000

To examine the role that each N domain plays in phase separation with RNA, we incubated fluorescently labeled g1-1000 with both nucleocapsid domains, as well as FL-N, and collected fluorescence and brightfield images. We found that both the FL-N and the CTD phase separated with g1-1000 at 37°C (Fig. 1i), while the NTD did not. To further explore the phase space, we prepared samples of each protein with g1-1000 at varied pH, temperature, and concentrations (Supp. Fig. S1). FL-N appeared sensitive to variations in conditions, forming droplets more readily at high temperatures, neutral pH, and a higher ratio of FL-N:g1-1000 (3.6 μM FL-N and 50 nM g1-1000 vs 9 μM FL-N and 50 nM g1-1000) (Supp. Fig. S1). In contrast, the NTD and CTD did not vary in their behavior – the CTD formed droplets under all tested conditions, while the NTD did not form droplets under any condition, leading us to conclude that the NTD does not play a major role in phase separation.

### Designing 14mer RNAs as models of secondary structure

Owing to the variety of structures in the g1-1000 RNA (i.e. Fig. 1e, Supp. Fig. S2), it is difficult to ascertain molecular-level details of where on the viral RNA the nucleocapsid binds. Analysis of RNP-MaP data^10^, which report on where N and the viral genomic RNA interact, show that the number of unpaired bases in a segment of g1-1000 correlate with N binding, suggesting that N binds preferentially to ssRNA (Fig. 2a). To validate this preference, we synthesized a 14- nucleotide RNA fragment (ss-14mer, Fig. 2d) along with its reverse complement to build a strong double-stranded RNA (ds-14mer, Fig. 2d). We designed 14-mer RNAs with a GC-rich sequence selected from short RNA sequences that were computationally tested for low self-pairing propensity for both the sequence and its reverse complement, and therefore ensure structural and conformational homogeneity. The single stranded structures of ss-14mer and its reverse complement were confirmed with RNAfold^41^ (Dot plots in Supp. Fig. S3a). RNAcofold predictions of free energy indicated that ds-14mer heterodimer is much stronger (-27 kcal/mol) than homodimer structures (-8.7, -7.2 kcal/mol). Lastly, ^1^H NMR spectra confirmed each structure of the RNAs. Imino protons in paired RNA bases have a characteristic ^1^H chemical shift above 11 ppm, allowing for experimental assessment of dsRNA content (Fig. 2d). The absence of peaks for the ss-14mer in this region is consistent with absence of any paired structure (Fig. 2d). Upon annealing with the reverse complement sequence, the fingerprint of ^1^H peaks in the 5-6 ppm region changes (Supp. Fig. S3c,d) and new peaks characteristic of paired imino protons appear at 11-15 ppm, confirming the formation of a double-stranded RNA (Fig. 2d).

**Figure 2:**
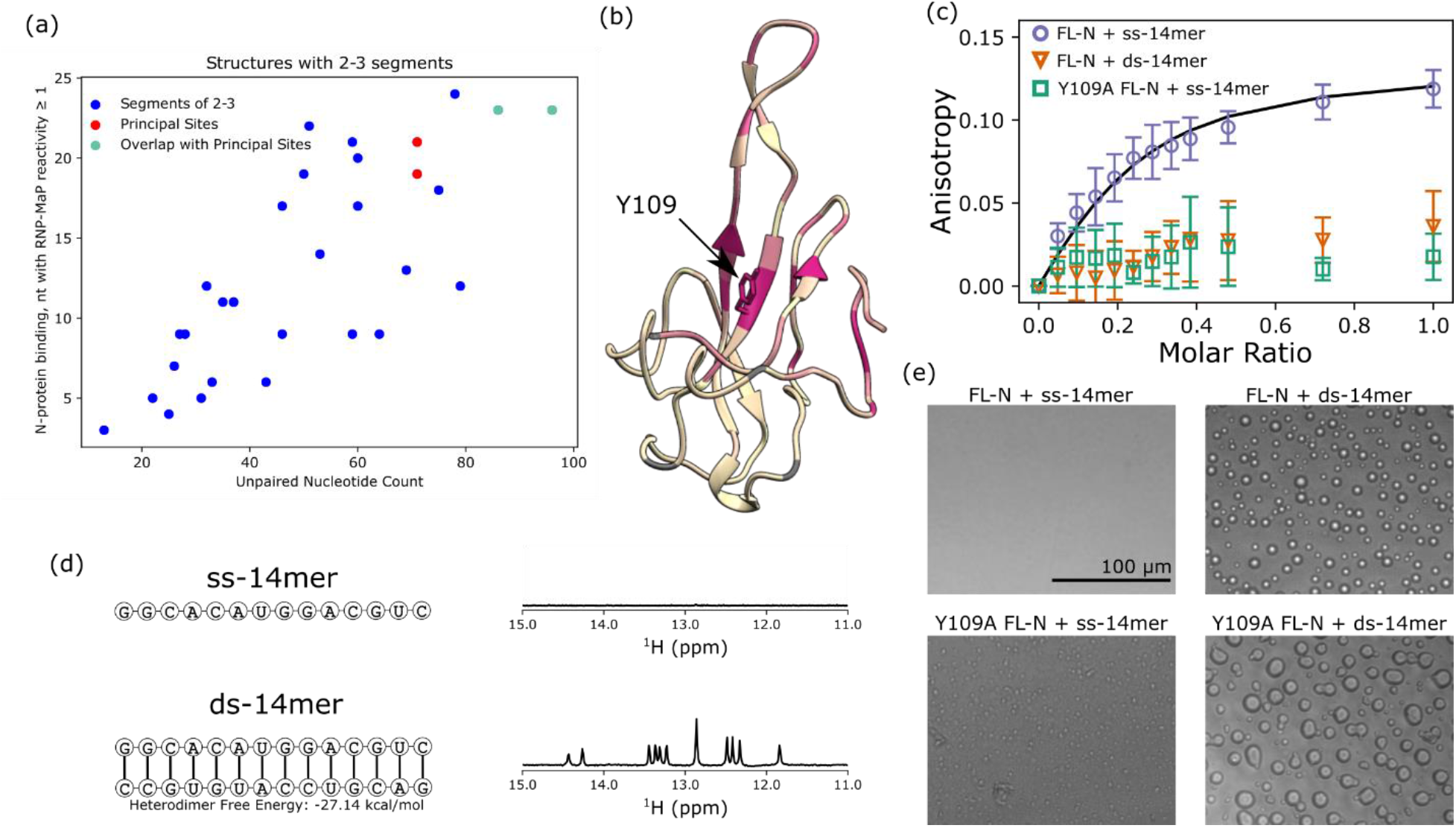
Overview of 14mer RNA interactions with FL-N and Y109A. (a) Plot of number of unpaired bases in structures consisting of 2-3 segments (see Methods) within the g1-1000 RNA against the RNP-MaP reactivity of the region^10^. The segments containing the principal sites as identified by Iserman et. al.(2020)^10^ are colored red, and those that overlap it are colored green. (b) Ribbon diagram of the NTD (PDB 7CDZ), colored by chemical shift perturbations induced in the NMR spectrum of the NTD by the Y109A mutation. Color is scaled from cream to red mapped to 0 to 0.2 CSP. See Supp. Fig. S5 for a plot of these perturbations and spectrum of the mutant NTD. (c) Fluorescence anisotropy titrations of FL-N or Y109A FL-N into fluorescently labeled ss- 14mer (purple, green) or ds-14mer (orange) at 50 nM. Uncertainty is taken from the standard deviation of three experimental replicates. Fit with a locked n=4 stoichiometry is shown in black for FL-N with the ss-14mer. (d) Sequence and structure of ss-14mer (top) and ds-14mer (bottom) used in this study, next to ^1^H NMR spectra in the 11-15 ppm region for each RNA. Peaks in this region are present for the ds-14mer (bottom) but not the ss-14mer (top) indicating base pairing. (e) Brightfield images of FL-N or Y109A FL-N mixed with the ss-14mer and ds-14mer, at FL-N concentration of 3.6 μM and RNA concentration of 18 μM.

### The NTD in FL-N contains the primary RNA binding site with preference to ssRNA

We next investigated binding between FL-N and the 14mer RNAs by fluorescence anisotropy. From titrations of FL-N into fluorescently labeled 14mer RNAs, we find that while FL-N binds tightly to the single-stranded RNA, it binds only weakly to the ds-14mer, inducing little change in anisotropy at the concentrations measured (Fig. 2c). In comparison, the anisotropy of the ss- 14mer rises quickly and nears a plateau even at sub-stoichiometric ratios of FL-N to RNA, suggesting a high stoichiometry of binding. Indeed, in our attempts to fit the anisotropy curve to a model of binding, we found that a stoichiometric ratio of at least 4:1 RNA:FL-N was necessary to fit the curve (Supp. Fig. S4), corresponding to a Kd of 80 nM. While this fit cannot be interpreted as an accurate measurement of the affinity between FL-N and the ss-14mer, it provides an estimate that FL-N binds the ss-14mer RNA with an affinity on the nanomolar scale. While similarly uncertain, the high stoichiometry is consistent with the fact that that the NTD, CTD, and disordered linker all bind RNA^13, 15, 20, 21^.

To characterize the role of the NTD in the nucleocapsid’s binding preference to ssRNA, we examined the same interaction using a Y109A mutant of FL-N. Y109 is in the RNA-binding groove of the NTD (Fig. 2b), and the Y109A mutation abrogates RNA-NTD binding^10, 34^. No structural characterization of the Y109A NTD has yet been reported, however. To ensure the structural fidelity of the mutant, we examined the Y109A NTD by NMR and demonstrated that the mutation primarily induces chemical shift perturbations at residues near Y109 in structure (Fig. 2b). Further, measurements of ^1^H R1 and R2 spin-relaxation rates exhibited minimal difference between the WT and Y109A NTD (Supp. Fig. S5c), indicating minimal change in the dynamic behavior of the protein. With this evidence of the integrity of the mutant NTD structure, we performed the same titration of Y109A FL-N into the ss-14mer RNA. Interestingly, the mutation nearly eliminates the change in anisotropy we see with the WT (Fig. 2c), indicating that the NTD structure around 109 is the site on N that binds tightly to the ss-14mer.

To determine whether the two RNA structures show any difference in their propensity to phase separate with FL-N, we collected images of both FL-N and the Y109A mutant of FL-N with the ss-14mer and ds-14mer RNAs. While no droplets were observed with the ss-14mer and WT FL-N, large liquid-like droplets were obtained with the ds-14mer RNA (Fig. 2e), providing further evidence for the hypothesis that double stranded RNA is a driver of phase separation, as has been proposed in prior literature^34^. For the Y109A FL-N, however, the mixture surprisingly formed droplets with both the ss- and ds-14mer, indicating that abrogation of binding at the NTD primary site promotes phase separation with ssRNA.

### Titrations with the individual domains confirm the NTD’s preference for ssRNA over dsRNA

To further confirm the NTD as the source of strong binding to ss-RNA, we next examined binding of each domain in isolation to our 14mer RNAs. Fluorescence anisotropy experiments indicate that the ss-14mer binds most tightly to the NTD, and only weakly to the CTD and the Y109A NTD (Fig. 3a). Experiments with the ds-14mer produce similar results, showing that the NTD binds the ds-14mer more tightly than the Y109A NTD or the CTD. Binding curves fit to 1:1 stoichiometry for the NTD binding to the ss-14mer and ds-14mer give affinities of 22 and 30 μM, respectively (Fig. 3a,b), showing that at this concentration the NTD binds both RNA structures but appears to favor the ss-14mer. We additionally performed electrophoretic mobility shift assays (EMSA) on the NTD and Y109A NTD with the ss- and ds-14mer. EMSAs confirm that the NTD binds the ss- 14mer slightly more strongly than the ds-14mer (as is apparent at 50 μM NTD comparing Fig. 3c to 3e), and that the Y109A mutation further weakens interactions between the NTD and each RNA. EMSA experiments further demonstrated that the Y109A NTD binds the ds-14mer marginally weaker than the ss-14mer, a difference which was not observable by anisotropy due to the relative weakness of both interactions and the low concentrations used in the anisotropy experiments.

**Figure 3:**
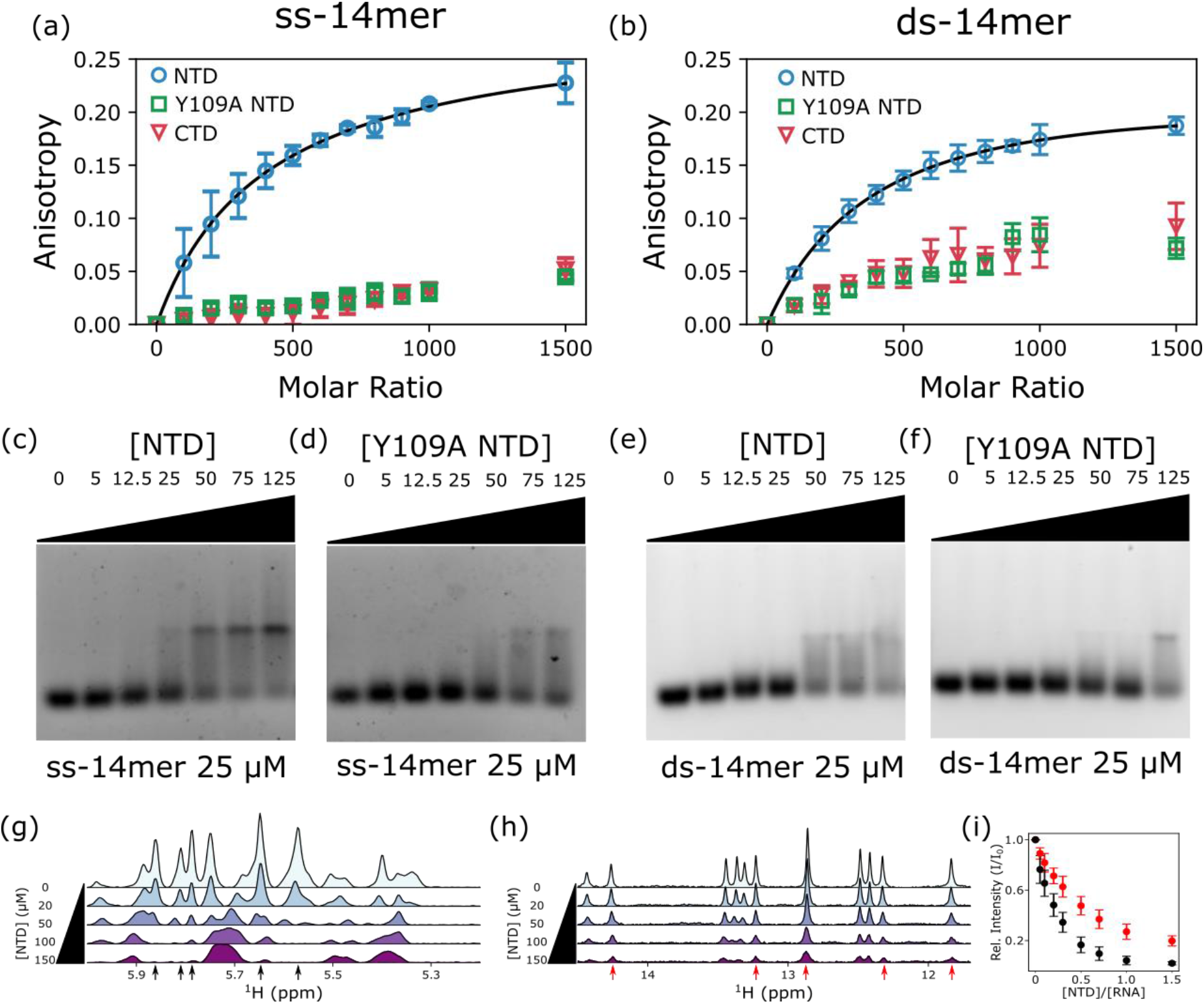
Nucleocapsid domain interactions with ss- and ds-14mer RNAs. (a,b) Fluorescence anisotropy titrations of the nucleocapsid domains into (a) the ss-14mer or (b) ds- 14mer at 50 nM. Fit is shown in black for the wt NTD in both panels. (c-f) EMSA of the NTD or Y109A NTD titrated into the ss-14mer or ds-14mer. RNA concentrations are fixed at 25 μM, while NTD concentrations are varied from 0 to 125 μM. (g) 1H NMR spectra in the 5.2-6 ppm region of 100 μM ss-14mer at increasing concentrations of the NTD up to 150 μM. (h) 1H NMR spectra in the 11 to 15 ppm region of 100 μM ds-14mer at increasing concentrations of the NTD up to 150 μM. The 5.2-6 ppm region for the ds-14mer is available in Supp. Fig. S6. (i) average peak intensities of the five peaks labeled by arrows in black for the ss-14mer (g), and red for the ds- 14mer (h).

To further examine the difference in binding affinity between ss-14mer and ds-14mer, we performed NMR titrations of the NTD into solutions of the RNAs (Fig. 3g,h). ^1^H NMR spectra of the ds-14mer are ideally suited for this analysis, due to the presence of imino proton peaks in the range of 11-15 ppm (Fig. 3h), outside the chemical shift region for protein amides. Although it is less isolated, the ^1^H spectra of the ss-14mer also contain a series of peaks where no proton resonances appear in the NTD, at 5-6 ppm (Fig. 3g). For both the ss-14mer and ds-14mer, addition of the NTD induced peak attenuation uniformly in every RNA peak. We selected 5 peaks with minimum overlap from each spectrum (black and red arrows in Fig. 3g,h) for analysis and plotted their average peak intensity ratio against the NTD concentration (Fig. 3i). We found a greater peak attenuation for the NTD with the ss-14mer, indicating tighter binding for the ss- 14mer, consistent with our EMSA and fluorescence anisotropy results.

### ssRNA and dsRNA bind the same charged groove on the NTD

Titrations of the ss-14mer RNA into ^15^N labeled samples of the NTD demonstrate that the RNA binds at the protein’s charged face. The addition of the ss-14mer induces multiple changes in the protein spectrum. Peaks for many residues either disappear due to peak broadening induced by intermediate exchange or shift due to a fast exchange with the RNA-bound state (Fig. 4a). The 29 disappearing peaks are for residues that cluster at the cup and form the RNA-binding face of the domain (Fig. 4b), including many residues in the 90-110 region, which corresponds to the central β hairpin of the cup. This indicates that the ss-14mer binds the NTD at that site, as demonstrated in previous work^13, 15^, broadening peaks where it binds. Chemical shift perturbation (CSP) plots indicate that residues 60-65, 128-130, and the C-terminus have the largest chemical shift perturbation (Fig. 4c). Mapping these to the structure (Fig. 4b) reveals that perturbations in the amide’s local environment are clustered at the periphery of the RNA-binding site, particularly at the side of the RNA-binding cup (e.g. D63, L167). This secondary patch of shifting peaks aligns well with results from Caruso et. al.(2022)^18^, which reported a similar patch including residues 64- 66 and 165-167 in titrations with 7-mer RNAs. While it is possible that these residues may correspond to an event distinct from binding at the charged face, the proximity of perturbed residues to the primary binding face leads us to conclude that all changes in the NTD spectrum are induced by the primary binding event. Therefore, whether peaks disappear or are shifted is dependent on the proximity of the residue to the binding face.

**Figure 4:**
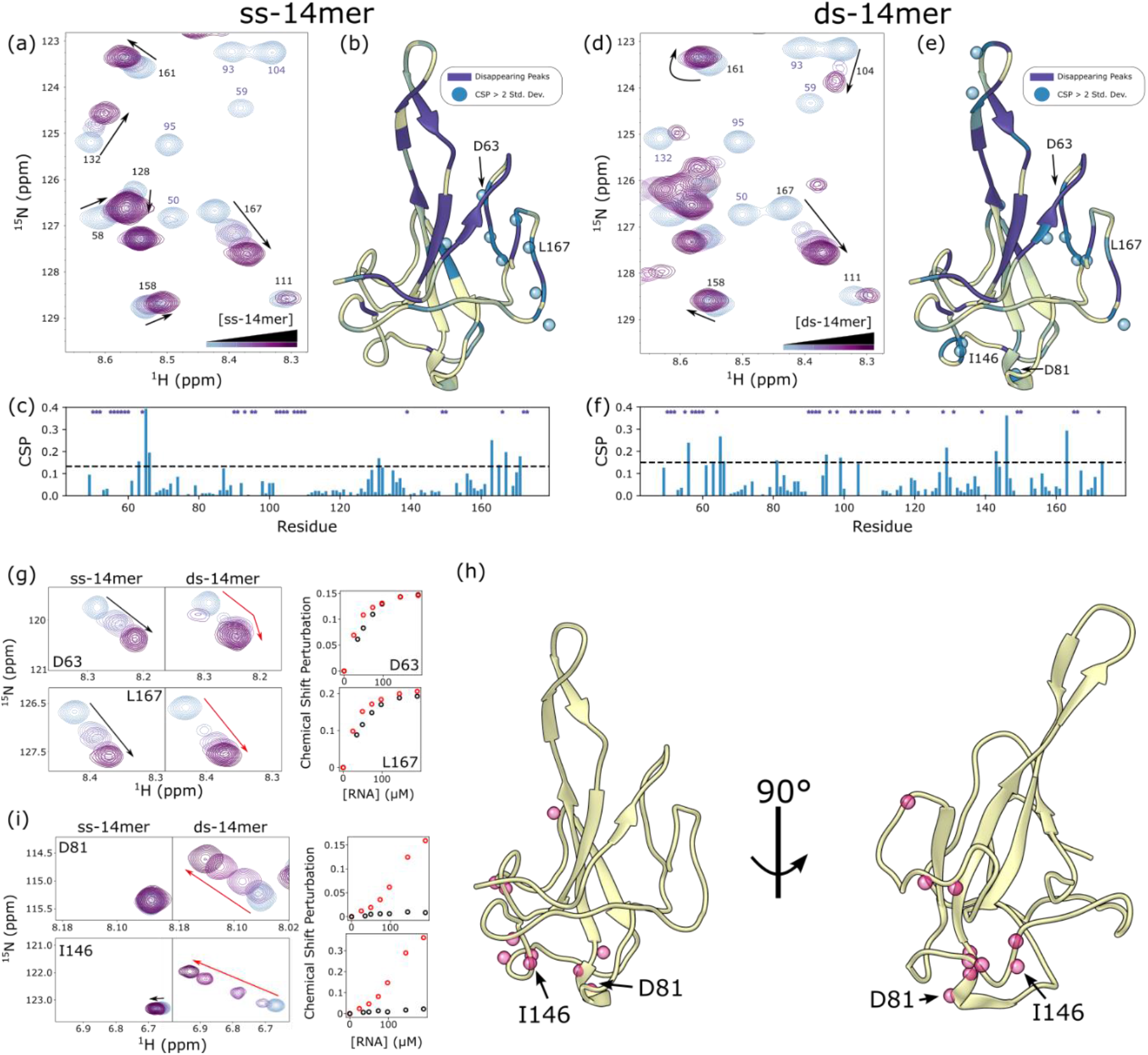
Titrations of 14mer RNAs into the NTD. (a,d) snapshot of ^1^H-^15^N HSQC spectra of the NTD (100 μM) at increasing concentrations of the ss-14mer (a) or ds-14mer (d), up to 200 μM RNA. Residues in a fast exchange regime are labeled in black, while those in an intermediate exchange regime are in purple. (b,e) ribbon diagram of the NTD (PDB 7CDZ) colored by their response to titrations of the ss-14mer (b) or ds-14mer. Residues that disappear due to intermediate exchange are colored purple, while those that shift are colored from cream to blue with increasing CSP. Residues with CSPs > 2 std. dev. are labeled with a blue sphere. (c,f) plot of CSPs for each titration (c for ss-14mer, f for ds-14mer) as a function of residue number. A black dashed line represents two standard deviations of CSPs, and residues whose peaks disappear due to intermediate exchange are labeled with a purple star at the top of the plot. (g,i) Window showing shifts of peaks D63 and L167 (g) or D81 and I146 (i) as a function of RNA concentration for both the ss- and ds-14mer. Plots to the right show CSPs as a function of RNA concentration, black for the ss-14mer and red for the ds-14mer. (h) Ribbon diagram of the NTD (PDB 7CDZ) with pink orbs representing all residues with linear CSP curves in response to the ds-14mer titration (see Supp. Fig. S6).

The ds-14mer induces a similar pattern of CSPs and peak disappearance (Fig. 4d), mapping to the same site as the ss-14mer (Fig. 4e). This similarity indicates that binding to the ds-14mer happens at the same face as to the ss-14mer. While this confirms that the ss-14mer and ds-14mer bind at the same site, some differences between the two RNA titrations are apparent (Fig. 4e,f). Rather than clustering together on the structure as in the ss-14mer titration, CSPs appear at several regions of the NTD. In addition to the cluster around D63 and L167, there are large shifts at the loop of the β hairpin, and an additional cluster of residues on the opposite face of the protein (e.g. I146 and D81).

### NMR identifies another binding site on the NTD

Chemical shift perturbations in the ds-14mer titration exhibit two distinct curves, indicative of two separate binding events. Residues such as D63 and L167 which are near the primary site exhibit behavior characteristic of tight binding – plots of the perturbation against RNA concentration level off asymptotically as the RNA concentration increases past a 1:1 ratio, indicating the NTD is near saturation at the primary site. This holds true for both the ss-14mer and ds-14mer titration at D63 and L167 (Fig. 4g), and other fast-exchanging residues at the same location in the structure. However, equivalent plots for D81 and I163 in the ds-14mer titration reveal a qualitatively different behavior (Fig. 4h). In these cases, the CSP increases roughly linearly with the RNA concentration, indicative of weak, nonspecific binding. Intriguingly, this behavior appears in only the ds-14mer titration, as the same peaks do not move at all in the ss-14mer titration (Fig. 4h). Analysis of the CSP plots for each residue in the ds-14mer titration with this distinction in mind reveals a cluster of residues around D81 and I146, including the helix at residues 79-82, residues 119 and 120, and residues 143-146 (Fig. 4i) (Supp. Fig. S6). As the CSPs at this site differ substantially from those near the primary binding site, we conclude that this shift corresponds to a second, weak binding surface of the NTD.

### Binding at the additional site is observed in the Y109A mutant

To further explore this second site and demonstrate that it is not an artefact of the size of the ds- 14mer, we investigated binding of the Y109A NTD to the 14mer RNAs. While the Y109A NTD binds to RNA only weakly, as seen in the anisotropy and EMSA experiments (Fig. 3), NMR conditions, which necessitate high sample concentrations, can still inform on the site of weak interactions. For both the ss- and ds-14mer, the RNAs induce a similar mix of peak behavior in the Y109A NTD as in the WT (Fig. 5a,b). The ss-14mer induces a similar pattern of disappearing residues in the mutant as in the WT NTD. Interestingly the ds-14mer titration results in the disappearance of fewer peaks (only 16, compared to 32 in the WT NTD), which we take as evidence of difference in exchange regime, consistent with the Y109A ds-14mer interaction being the weakest of those tested. CSPs for fast-exchanging residues occur in a similar pattern to the WT NTD (Fig. 5a,b, Supp. Fig. S7c), which, taken with the disappearance of the same peaks for residues at the primary binding site, indicates that the Y109A NTD binds both ss- and ds-RNA at the same site as the WT NTD. Titrations of the NTD with RNA (as in Fig. 3g,h) confirm the general trends of weaker binding observed by anisotropy and EMSA experiments (Supp. Fig. S7).

**Figure 5:**
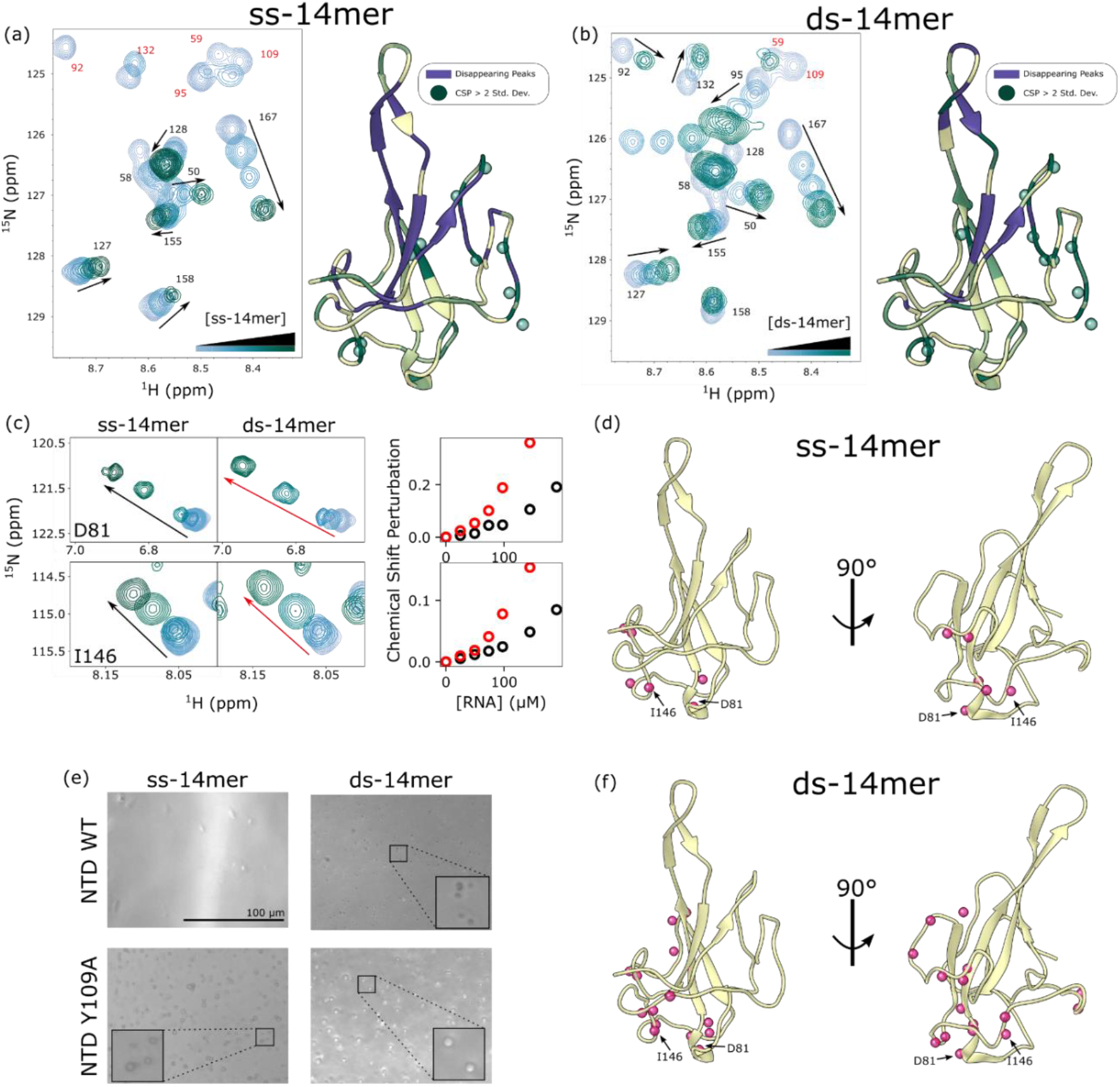
Titrations of 14mer RNAs with Y109A NTD. (a,b) (left) snapshot of ^1^H-^15^N HSQC spectra of the Y019A NTD (100 μM) at increasing concentrations of the ss-14mer (a) or ds-14mer (d), up to 200 μM RNA. Residues in fast exchange regime are labeled in black, while those in intermediate exchange regime are labeled in purple. (right) ribbon diagram of the NTD (PDB 7CDZ) colored by their response to titrations of the ss-14mer (b) or ds-14mer (e). Residues that disappear due to intermediate exchange are colored purple, while those that shift are colored from cream to green with increasing CSP. Residues with CSPs > 2 std. dev. are labeled with a green sphere. (c) Window showing shifts of peaks D81 and I146 as a function of RNA concentration for both the ss- and ds-14mer. Plots to the right show chemical shift perturbation as a function of RNA concentration, black for the ss-14mer and red for the ds-14mer. (d,f) Ribbon diagram of the NTD (PDB 7CDZ) with pink orbs representing all residues that exhibit a linear CSP curve with the ss-14mer (d) or the ds-14mer (f) (see Supp. Fig. S7 for all CSP profiles). (e) Brightfield images of the NTD or Y109A NTD at 100 μM mixed with the ss-14mer or ds-14mer at 200 μM collected at 20 mM NaCl and 37 °C.

Interestingly, the cluster of residues with high CSPs map to the same surface opposite the primary binding site with both ss and ds-RNA. Revisiting D81 and I146 peak changes, we see large perturbations in chemical shifts with both titrations (Fig. 5c). While the ds-14mer induces the greatest shift, the interaction is clearly present with the ss-14mer titration as well, and plots of CSPs as a function of RNA concentration exhibit the same linear-like behavior seen in the WT NTD interaction with the ds-14mer (Fig. 5c). Only 6 residues exhibit this behavior in the ss-14mer titration (Fig. 5d), while 16 can be observed with the ds-14mer (Fig. 5f), including several in the 120-130 loop. The fact that both the ss-14mer and ds-14mer induce the same chemical shift perturbations in the mutant, demonstrate that this secondary site is not specific to the double-stranded RNA structure. Notably, the number of shifting peaks, as well as the magnitudes of the shifts observed, correlate inversely with the tightness of binding as observed by anisotropy and EMSA assays. For example, the ss-14mer-NTD interaction has the highest affinity and shows no evidence of binding at this secondary site, while the ds-14mer-Y109A NTD interaction is the weakest and shows the most evidence of this secondary interaction.

At NMR conditions, the propensity of the NTD to form liquid droplets is modulated both by RNA structure and mutation in the NTD. During our NMR experiments, we noticed that the Y109A mutant phase separated with RNA under some conditions. In fact, we were unable to collect a spectrum of the mutant NTD with ds-14mer above 150 μM RNA due to the sample’s propensity to form droplets (see methods for details). To examine this phenomenon more comprehensively, we collected images of samples of the WT and mutant NTD with the ss- and ds-14mer at NMR conditions. At 37 °C, a temperature that promotes nucleocapsid phase separation relative to the 25 °C used in the NMR experiments, the NTD forms droplets with the ds-14mer (Fig. 5e). Further, the Y109A mutation promotes droplet formation, forming large droplets with both the ss- and ds- 14mer. At 25 °C, only the ds-14mer and Y109A NTD formed droplets (Supp. Fig. S8). Interestingly, this pattern matches what we see for FL-N as well (Fig. 2e): the WT protein will only phase separate with the ds-14mer, while the Y109A mutant will phase separate with either RNA.

### The CTD shows preference for binding dsRNA at its alpha face

We last turned our attention to interactions between the CTD and the 14mer RNAs. Attempts to assay this interaction by NMR were hampered overall by peak broadening, minimal perturbations in chemical shifts, and the propensity of the CTD to phase separate with RNA, which necessitated collecting spectra at high salt conditions. Addition of g1-1000 to the CTD produced no chemical shift perturbations, but induced dips in peak intensity at parts of the CTD corresponding to the domain’s binding face (Fig. 1g). Titration of the 14mer RNAs into the CTD produced similar behavior, decreasing peak intensities but not inducing chemical shift perturbations. For example, spectra at 0.66:1 RNA:CTD show weak peaks with no changes in chemical shift for most residues (Fig. 6a). While relative peak intensities at this concentration ratio are roughly uniform across the protein sequence when bound to the ss-14mer (Fig. 6b), the ds-14mer-bound spectrum is highly heterogeneous (Fig. 6c). Peak intensities are weak, and many peaks are missing from the N-terminus of the CTD, as well as for residues 300-320, both of which are on the α face of the CTD, suggesting this drop in intensity is due to interaction with RNA at this charged face (Fig. 6e). It is intriguing, however, that this significant intensity drop appears to occur only with the ds-14mer, although both RNAs clearly bind to the CTD. The difference between spectra may indicate a difference in the two RNA-CTD interactions, suggesting the ds-14mer may interact with the α face, while the ss-14mer binds nonspecifically, although other factors such as the rate of exchange between the apo and bound state may also impact peak intensities.

**Figure 6:**
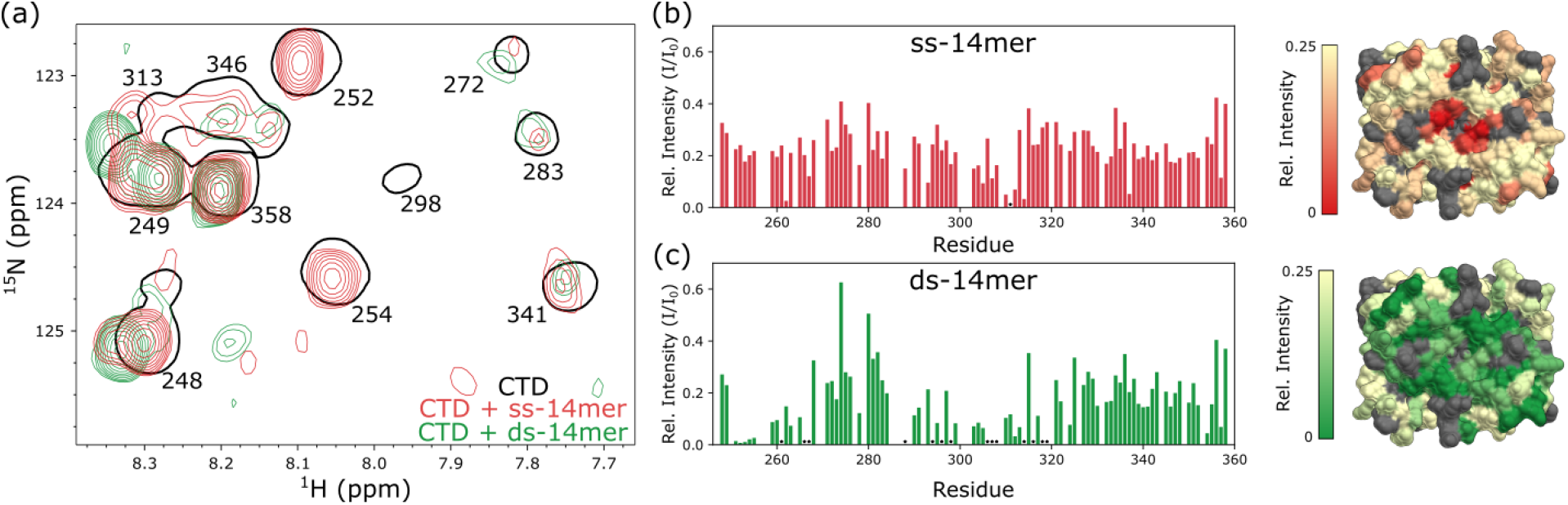
Binding between the CTD and 14mer RNAs. (a) Snapshot of the CTD spectrum bound to the ss-14mer (red) and ds-14mer (green) at 66 μM 14mer RNA and 100 μM CTD. Outlines of the apo spectrum are shown in black, along with the assignments for each residue. (b,c) Relative peak intensity (to the apo spectrum) plotted (left) and mapped to the protein surface (right) for the ss-14mer (b) and ds-14mer (c) bound CTD. Peaks that were not detectable are marked with a star. Ribbon (left) and surface (right) diagram of the CTD colored by the relative peak intensity of the bound spectrum. For the protein surfaces, white represents a relative intensity of 0.25 or greater, while red (d) or green (e) represents an intensity of 0. Unassigned or overlapped peaks are colored gray.

## Discussion

The SARS-CoV-2 nucleocapsid protein interacts multivalently with RNA to facilitate a range of functions within an infected cell. Due to the complexity and multiplicity of domains of both the nucleocapsid and the viral RNA, there is a large variety in the potential structures of nucleocapsid-RNA complexes. To identify structural preferences within the viral genomic RNA, we designed 14mer RNAs consisting entirely of ssRNA (ss-14mer) as a model of regions of unpaired RNA and dsRNA (ds-14mer) as a model for paired stem regions and examined interactions between these RNAs and both nucleocapsid domains, as well as the full-length protein, and their propensity to form liquid condensates. We find that both domains bind to both RNAs, each with different affinities and different propensities for phase separation. In particular, interactions between N and ssRNA are strong, and a mutation that abrogates binding at the NTD dramatically weakens binding between N and RNA, indicating that high-affinity interactions are mediated through single-stranded RNAs and the primary site of the NTD. We additionally describe a second RNA-binding face of the NTD that interacts only weakly with RNA and confirm the α face of the CTD as the region that binds RNA, with preference for dsRNA. Lastly, we examine how these interactions correlate with the ability of the nucleocapsid to phase separate with RNA, demonstrating that multiple weak, nonspecific interactions at the CTD and at the NTD’s secondary face drive liquid droplet formation.

### Two faces of the NTD bind RNA

In addition to confirming the primary binding face, NMR titration experiments identified a second site containing several residues (e.g. D81, I146 in Fig. 4, Fig. 5) that exhibit behavior distinct from those at the primary site. As noted above, the degree of shift for this cluster correlates inversely with binding affinity – the tight-binding NTD-ssRNA complex shows no shifts at the secondary site, while titration of dsRNA into the Y109A NTD induces the greatest shifts (with the other two complexes in-between). One possible explanation for this observation is that the NTD-RNA interaction is in equilibrium between two mutually exclusive bound states. As such, when the primary interaction weakens, the relative favorability of the secondary interaction is increased.

The degree of binding at the second site also correlates with the propensity of the NTD to form liquid droplets. For example, the only complex that did not form droplets at 37°C was the WT NTD with ssRNA, while the Y109A NTD formed droplets most readily with dsRNA, even at lower temperatures (Supp. Fig. S8). Given that experimental and computational models of phase separation emphasize the importance of weak, multivalent interactions^42, 43^, we propose that mutations in the NTD that drive binding at the secondary site allow the NTD to multivalently bridge several RNAs, forming the complex interaction network required for phase separation.

### The nucleocapsid in context

Several studies have now suggested that the NTD may exhibit preference for some RNA structures over others^18, 25, 29^. Most recently, work from Korn et al^29^ suggested that the NTD exhibits a slight preference for single-stranded RNA structures, based on EMSA experiments. Here we confirm through several methods that the NTD binds preferentially to ssRNA over dsRNA. While the preference for ssRNA is weak, our anisotropy data on N indicate that the NTD’s preference for ss-RNA is greater in the full length protein, suggesting that either multivalency or the disordered linker may play a role in RNA structure discrimination. The observation that a single mutation in the NTD significantly reduces binding in the full length protein, however, argues for a more prominent role of the NTD. Nonetheless, how other regions of N outside of the NTD impact RNA structure and motif discrimination remain a major question going forward.

The formation of N-RNA condensates with various RNA structures demonstrate that N has a remarkable propensity to form liquid-liquid phase separated droplets with RNA. Several studies have suggested that the nucleocapsid’s CTD drives condensate formation^34, 44,45,46^, modulated by the linker connecting the two domains, which promotes the formation of liquid-like condensates, especially when phosphorylated^12, 24, 47, 48^. Peptides designed to interrupt CTD dimerization also interrupt droplet formation^49^, suggesting that multivalency of the dimeric N promotes separation as well. We demonstrate that the CTD strongly drives phase separation, consistently condensing with g1-1000 at all conditions we tested. In fact, in our hands, N has a reduced propensity to form liquid droplets with genomic RNA relative to the CTD at equivalent concentrations, suggesting that interactions at the NTD or IDRs may inhibit condensation. Consistent with this hypothesis, we find that mutation of the NTD to abrogate RNA binding promotes phase separation (Fig. 2e, Fig. 5e), possibly through binding at a secondary face on the NTD. It bears mentioning that this conflicts somewhat with prior work on the Y109A mutant, which has indicated that the mutation either inhibits phase separation^34^ or alters droplet morphology^34^. One potential explanation for this lies in the difference in RNA length between prior studies which used RNAs of 1000 nt in length and our work focused on 14mers. A recent study^34^ has demonstrated that mutating viral genomic RNA to increase its double-stranded content promotes phase separation, consistent with our findings that both FL-N and the NTD phase separate more readily with dsRNA than ssRNA. Mechanistically, these observations suggest that weak, nonspecific binding interactions - binding between CTD and RNA or binding at the NTD’s secondary face - promote separation while strong interactions, such as between the NTD and ssRNA, inhibit it. The study presented here firmly establishes this is the case.

### Conclusions and Outlook

Like many viral proteins, the SARS-CoV-2 nucleocapsid protein plays multiple roles in viral particles and within infected cells. These roles primarily revolve around the organization and protection of viral RNA, along with facilitation of viral transcription and replication. N-RNA complexes form a cornerstone of the virus functions, and their interactions are dependent on variation in RNA sequence and structure. Binding between the nucleocapsid and model RNA structures as extensively characterized here demonstrate that the strongest N-RNA interaction independent of the RNA sequence is mediated by the NTD, which binds tightly to ssRNA, although it can bind dsRNA less strongly at the same site (Fig. 7a). Similarly, the CTD can bind either of the RNA structures but with weaker affinity in comparison to the NTD, occurring only at high concentrations of N and with some preference for dsRNA (Fig. 7b). For the full-length nucleocapsid, the NTD-ssRNA interaction outcompetes other interactions (Fig. 7c), although mutations that damage the NTD’s primary site abolishes viral RNA type discrimination^29^ by altering the equilibrium between binding at the primary face, at the NTD’s secondary face, and at the CTD (Fig. 7d). As the NTD-dsRNA interaction is weaker, the interaction is likewise in equilibrium with binding at the secondary face or the CTD (Fig. 7e). Liquid-liquid phase separated droplet formation of various N/RNA complexes demonstrate that multiple weak interactions promote droplet formation, while strong, specific interactions do not. The question of how the nucleocapsid is recruited to the sites of viral motifs cannot solely be answered by RNA secondary structure, however, and the question of motif sequence specificity remains an important one in the study of the nucleocapsid. With a firm understanding of how secondary structure relates to both specific and promiscuous interactions, future work investigating binding between the nucleocapsid and viral motifs will be well-equipped to explore how N senses and acts on viral RNA.

**Figure 7:**
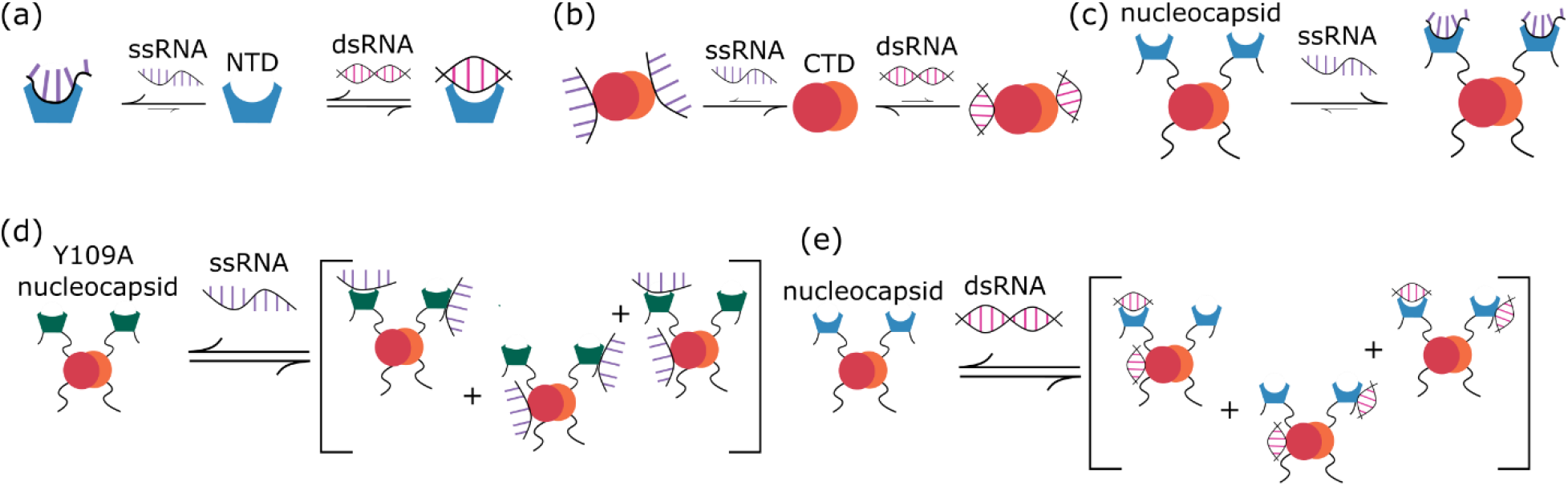
Overview of N-RNA interactions. Diagram of the NTD (a) and the CTD (b) binding to ss- and dsRNA. (c) Proposed model of FL-N tight binding to ssRNA through the NTD. (d) Proposed model of Y109A FL-N weak binding to ssRNA. Reduced affinity for ssRNA increases relative favorability of other interactions. (e) Proposed model of weak binding to dsRNA. As NTD- dsRNA binding is weaker, other interactions compete with NTD-RNA binding.

## Materials and Methods

### Protein expression and purification

FL-N, NTD, and CTD, as well as Y109A mutants of FL-N and NTD were expressed as described previously^13^. Both the NTD and CTD constructs were identical to those used previously^13^, covering residues 47-174 for the NTD and 231-361 for the CTD. Briefly, FL-N and Y109A FL-N were grown in 2x YT or MJ9 media to an OD600 of 0.6, then induced with 1 mM isopropylthio-beta-galactoside (IPTG) at 18°C for 20 hours. CTD was grown to an OD600 of 2 in terrific broth, then induced with 0.5 mM IPTG at 18°C overnight for unlabeled expression. For ^15^N labeling, cells of FL-N or CTD were grown in 2x YT to an OD of 0.7, pelleted at low speed, then washed and resuspended in ^15^N-enriched MJ9 media, grown for an hour then induced with 0.5 mM IPTG at 30°C for 4 hours (CTD) or 18°C overnight (FL-N). The NTD and Y109A NTD were expressed in either ZYM-5052 autoinduction media (for unlabeled protein) or MD-5052 autoinduction media with ^15^N ammonium chloride (for stable isotope labeling)^50^.

Proteins were purified using TALON His-tag purification protocol (Clonetech). Cells were lysed in lysis buffer (50 mM sodium phosphate buffer, 1 M NaCl, 1 mM NaN3, 5 mM Imidazole, pH 8.0, 0.6 mg/mL lysozyme with protease inhibitor) for 1hr at 4°C. Cells were further sonicated and centrifuged at 20000 RPM for 45 min to separate cell debris. The supernatant was incubated with Cobalt resin in a gravity column. All proteins were washed on-column with 3M NaCl (20 column volumes) to remove bound bacterial RNA. FL-N, Y109A FL-N, and CTD were eluted with 4 column volumes of elution buffer (50 mM sodium phosphate at pH 8.0, 300 mM NaCl, 1 mM NaN3, 350 mM Imidazole). Purified NTD and Y109A NTD were eluted by on-column proteolysis with 30 nM untagged fast-acting protease bdSENP1 for 1 hr at 4°C. Proteins were further purified on a Superdex 75 gel filtration column (GE Health) in 50 mM sodium phosphate, 150 mM NaCl, pH 6.5. The purity of the recombinant proteins, assessed by SDS-PAGE, was >95%. Protein concentrations were determined using absorbance at 280 nm, calculated from standard extinction coefficients of 1490 and 5500 M^-1^ cm^-1^ for tyrosine and tryptophan respectively. Purified proteins were either stored at 4°C and used within one week or flash frozen to -80°C for long term storage.

### RNA preparation

The ss-14mer (5’-GGCACAUGGACGUC-3’), either labeled with 5’ 6-carboxyfluorescein or unlabeled and reverse complement 14mer (5’-GACGUCCAUGUGCC-3’) were purchased from GenScript. The dsRNA was prepared by annealing the ss-14mer (5’-GGCACAUGGACGUC-3’) with the reverse complement (5’-GACGUCCAUGUGCC-3’) at equal concentration of each oligonucleotide and 50mM NaCl. The mixture was incubated at 60°C for 15 min and cooled slowly to 26°C^15^. The purity and size of ssRNA and dsRNA were confirmed by agarose gel. The in vitro transcription and purification of the g1-1000 RNA followed protocols described previously^13^.

### Analysis of RNP-MaP data

For analysis of the published RNP-MaP data in Fig. 2a, we divided the 5′ genome into structural regions consisting of 2-3 segments, as was observed with the structural regions of the principal binding sites. A segment is defined as a region of adjacent base pairs that can be interrupted by bulges or internal loops, but not multiloops or hairpin loops^10^. For each region, we calculated the unpaired nucleotide count and the N-protein binding in terms of the number of positions within the region that had an RNP-MaP reactivity greater than or equal to 1. It has been suggested that RNP-MaP experiments are biased towards regions of unpaired bases, which would offer an alternate explanation for the results presented here (Fig. 2a). However, recent analysis of a large number of RNP-MaP experiments^51^ appears to indicate that the technique has negligible preference for RNA secondary structre (Supp. Fig. S2 in Weidmann et al. (2021)^51^).

### NMR spectroscopy

NMR spectra of the NTD, Y109A NTD and CTD were collected on a Bruker 800 MHz Avance II HD spectrometer equipped with a triple resonance cryogenic probe. All NMR samples contained 7.5% D2O for signal locking, 1 mM sodium trimethylsilylpropanesulfonate (DSS) as a reference, and a commercial protease inhibitor cocktail (Roche applied Science). Samples used in RNA titrations additionally contained a Murine RNAse inhibitor (NE Biolabs) at a concentration of 1 unit/μL. Assignments for the NTD (BMRB 34511) and CTD (BMRB 50518) were transferred from published assignments^15, 52^. Assignments for the Y109A NTD were transferred from published assignments for the WT NTD, aided by a 3D NOESY-HSQC spectrum used to confirm assignments for residues with mutation-induced chemical shift perturbations. The mutant assignments are available at BMRB 52010. All spectra were processed using NMRPipe^53^, assignments were transferred and made in CCPNMR Analysis^54^, and titrations were analyzed in NMRViewJ^55^. One-dimensional ^1^H NMR spectra of RNA were analyzed using in-house python scripts utilizing NMRGlue^56^, and baseline-corrected with the doubly reweighted penalized least squares algorithm in the pybaselines library^57^. Chemical shift perturbations for all experiments were calculated as 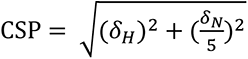, where *δ_H_* is the change in proton chemical shift and *δ_N_* is the change in nitrogen chemical shift, which is scaled by a factor of 1/5^58^.

Spectra of the NTD and Y109A NTD were collected at 25°C in a buffer of 20 mM sodium phosphate and 20 mM NaCl, at pH 6.5. For titrations of the ss- and ds-14mer, we collected spectra at 100 μM NTD titrated with RNA (concentrated to 4 mM to minimize dilution) up to 2:1 RNA:NTD, at intervals of 0.25,0.5,0.75,1,1.5 and 2:1 RNA:NTD. Figures of the spectra (Fig. 4a,d, Fig. 5a,b) show the 0, 0.25, 0.5, 1, 1.5, and 2:1 RNA:NTD titration points, with the exception of Fig. 5b, which is missing the 2:1 titration point due to phase separation at this ratio. During experiments, we found that some samples of NTD with RNA became cloudy at 37° C, an effect that was completely reversible upon incubation at 4°C. Images of these samples revealed this cloudiness to be the formation of liquid-liquid phase separated droplets (see Supp. Fig. S8). At 25°C only the 1.5 and 2:1 ds-14mer:Y109A NTD sample phase separated, indicating that other titrations are not impacted by phase separation. For the ds-14mer:Y109A NTD titration, we stopped titrating RNA at 1.5:1 to avoid having phase separation impacting protein concentrations.

For titrations of NTD into RNA, we performed experiments in a 3 mm NMR tube at 100 μM RNA in NTD NMR buffer and titrated NTD at 400 μM into the RNA samples up to a 1.5:1 NTD:RNA ratio, at intervals of 0.05, 0.1, 0.2, 0.3, 0.5, 0.7, 1 and 1.5:1 NTD:RNA. Plots (Fig. 3g,h, Supp. Fig. S7), show the spectra at 0, 0.2, 0.5, 1, and 1.5:1 NTD:RNA. Peak intensities from these titrations for plots in Fig. 3i and Supp. Fig. S7b were extracted using in-house python scripts utilizing NMRGlue. Peak heights were determined at each titration point by finding a local maximum in a window around the center of the unbound peak, to allow for small changes in chemical shift across the titration.

Experiments on the CTD were carried out at 25°C in a buffer of 20 mM sodium phosphate and 200 mM NaCl at pH 6 in a shaped NMR tube. At 200 mM NaCl we observed no evidence of phase separation of the CTD with RNA. All spectra were collected at 100 μM CTD, and at 1 μM g1-1000 (Fig. 1g) or 66 μM 14mer RNA (Fig. 6).

Measurements of ^15^N R1, ^15^N R2 rates, and {^1^H}-^15^N Heteronuclear NOEs were performed in the NTD NMR buffer conditions described above at protein concentrations of 500 μM. Temperature compensated versions of published sequences^59^, were used for ^15^N R1 and ^15^N R2 measurements. Relaxation rates were determined by fitting data to a single exponential decay in NMRViewJ. For spin-relaxation measurements, uncertainty was determined using triplicate measurements of one point within the relaxation curve and propagated to the rate using the Monte Carlo method. {^1^H}-^15^N heteronuclear NOE values were measured using a version of the sequence in^59^, with a recycle delay of 8 s. NOE (I/I0) was calculated from intensities measured in NMRViewJ, and uncertainty propagated through the standard deviation of the noise floor of the spectra.

### Fluorescence Anisotropy

Fluorescence anisotropy measurements, using 14mer RNAs labeled with 6-carboxyfluorescein were performed on a FluoroMax-3 spectrophotometer (Horiba Scientific). The excitation wavelength was set to 492 nm and the emission wavelength to 516 nm. For all experiments, a buffer of 50 mM sodium phosphate at pH 6.5, 150 mM NaCl, 1 mM NaN3 was used. For titrations with the N domains (Fig. 3a,b), the concentration of RNA was set to 50 nM and protein was titrated up to 1.5 μM (up to a 1500:1 NTD:RNA ratio). Titrations of FL-N and Y109A FL-N into the 14mer RNAs (Fig. 2c) were done at 50 nM RNA and titrated with protein up to a 1:1 ratio. NTD titrations were fit in GraphPad Prism 9 using the one site-total binding model. The titration of FL-N into ss- 14mer was fit with in-house python scripts (see Supp. Fig. S4), using a standard quadratic binding model^60^ with a fixed binding stoichiometry correction.

### Microscopy

Fluorescence and brightfield microscopy images were taken on a Keyence BZ-X700/BZ-X710 microscope and a 384-well plate (Cellvis P384-1.5H-N); images were processed using BZ-x viewer and BZ-x analyzer software. For all samples except the NTD with 14mer RNAs, protein and RNA were mixed at different ratios (listed in figure captions, Fig. 1i, Fig. 3e, Supp. Fig. S1) in 20 mM Tris, 150 mM NaCl, 1 mM DTT, pH 7.5 droplet buffer in a total volume of 30 µl, and incubated at 37°C for at least one hour followed by imaging. For the samples of NTD with 14mer RNAs (Fig. 5e, Supp. Fig. S8), NMR conditions (20 mM sodium phosphate pH 6.5, 20 mM NaCl) were used, and samples were incubated at either 25°C or 37°C for at least one hour prior to imaging.

### Electrophoretic mobility shift assays (EMSA)

NTD (WT and Y109A in 20 mM Tris, 150 mM NaCl, 1 mM DTT, pH 7.5 droplet buffer) binding to the ss-14mer and ds-14mer was visualized by electrophoretic mobility shift assay in 2.5% agarose gels. Increasing concentrations of protein in the range of 0-125 μM were added to 25 μM RNA and incubated for 20 min at room temperature in a total reaction volume of 10 μl. 2 μL loading dye was added to the reaction before loading. RNA bands were stained with Midori Green Nucleic Acid staining solution (Bulldog Bio. Inc. Portsmouth, NH) and visualized using a Bio-Rad Gel Doc Image system.

## Acknowledgements and Funding

We acknowledge Richard Cooley for the helpful discussions, and Seth Pinckney and Gretchen Fujimura for assistance with protein expression and purification. This work was supported by the U.S. National Science Foundation EAGER grant MCB 2034446 to E.B. We additionally acknowledge the support of the Oregon State University NMR facility funded in part by the National Institute of Health, HEI grant 1S10OD018518 and by the M.J. Murdoch Charitable trust grant 2014162 to E.B.

