## Supplementary Info for "RNA structure and multiple weak interactions balance the interplay between RNA binding and phase separation of SARS-CoV-2 nucleocapsid"

### Supplemental Information

#### Contents:

**Figure S1:** Temperature, pH and concentration ratio phase dependence of the nucleocapsid.

**Figure S2:** Secondary Structure of g1-1000.

**Figure S3:** Characterization of 14mer RNAs.

**Figure S4:** Analysis of the fluorescence anisotropy curve of FL-N binding to the ss-14mer.

**Figure S5:** NMR assignments and dynamics of Y109A NTD.

**Figure S6 :** Chemical shift perturbations for all secondary-site residues from titration of ds-14mer into the NTD.

**Figure S7 :** NMR analyses of Y109A NTD interaction with RNA.

**Figure S8 :** Phase separation of the NTD at NMR conditions.

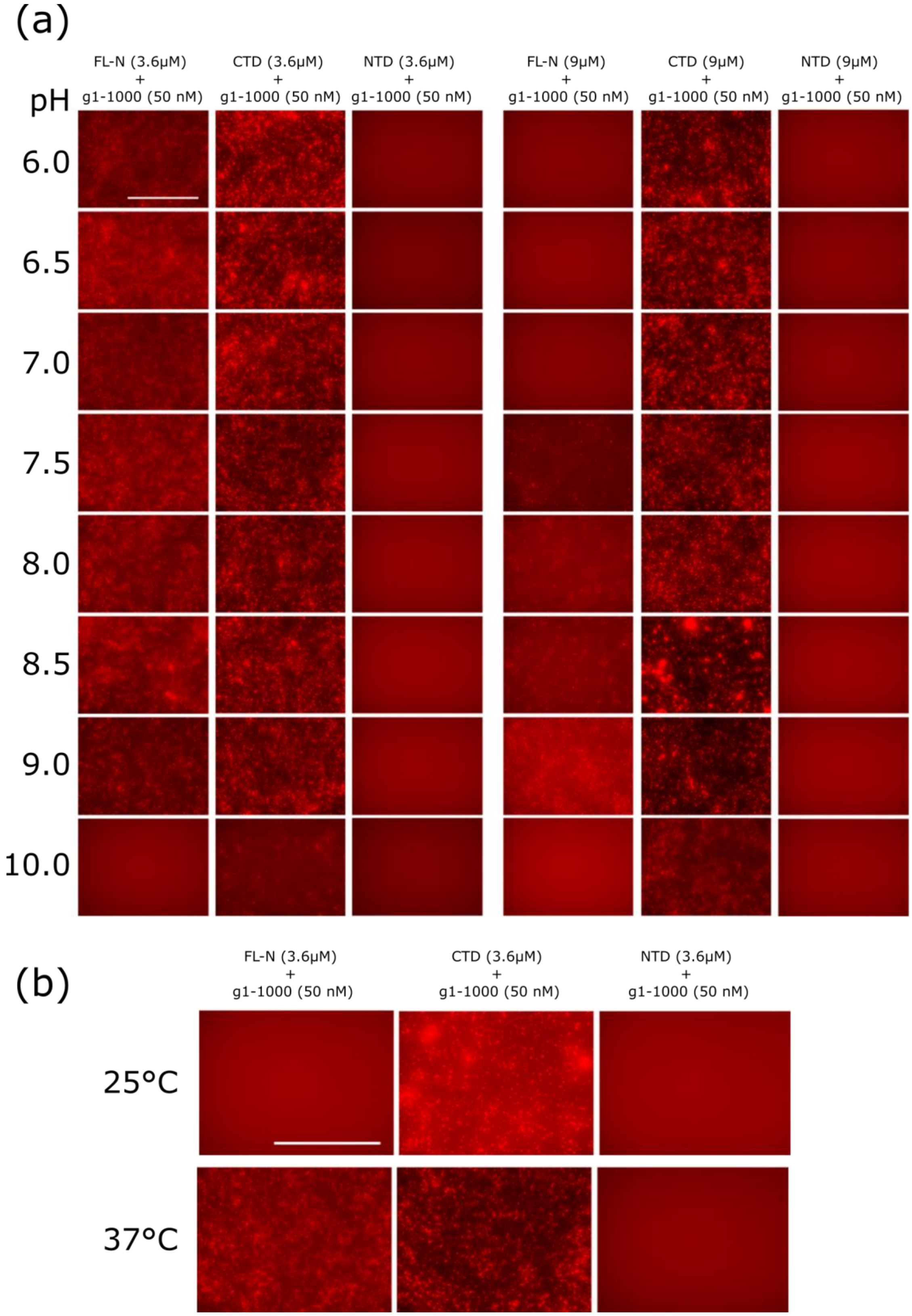


**Figure S1: Temperature, pH and concentration ratio phase dependence of the nucleocapsid.** Red fluorescence images of samples of FL-N, CTD and NTD with g1-1000 (buffer conditions of 20 mM Tris pH 7.5, 150 mM NaCl, 1 mM DTT). (a) 3.6 (left) or 9 (right) μM FL-N, CTD or NTD and 50 nM g1-1000, all images taken at 37°C. Scale bar in the top left panel is 200 μm and scale is the same for all images. (b) 3.6 μM FL-N, CTD or NTD and 50 nM g1-1000 at 25°C (top) and 37°C (botton). Samples in both panels were equilibrated at their experimental temperature for 90 minutes prior to imaging. Scale bar in the top left panel is 200 μm and scale is the same for all images.

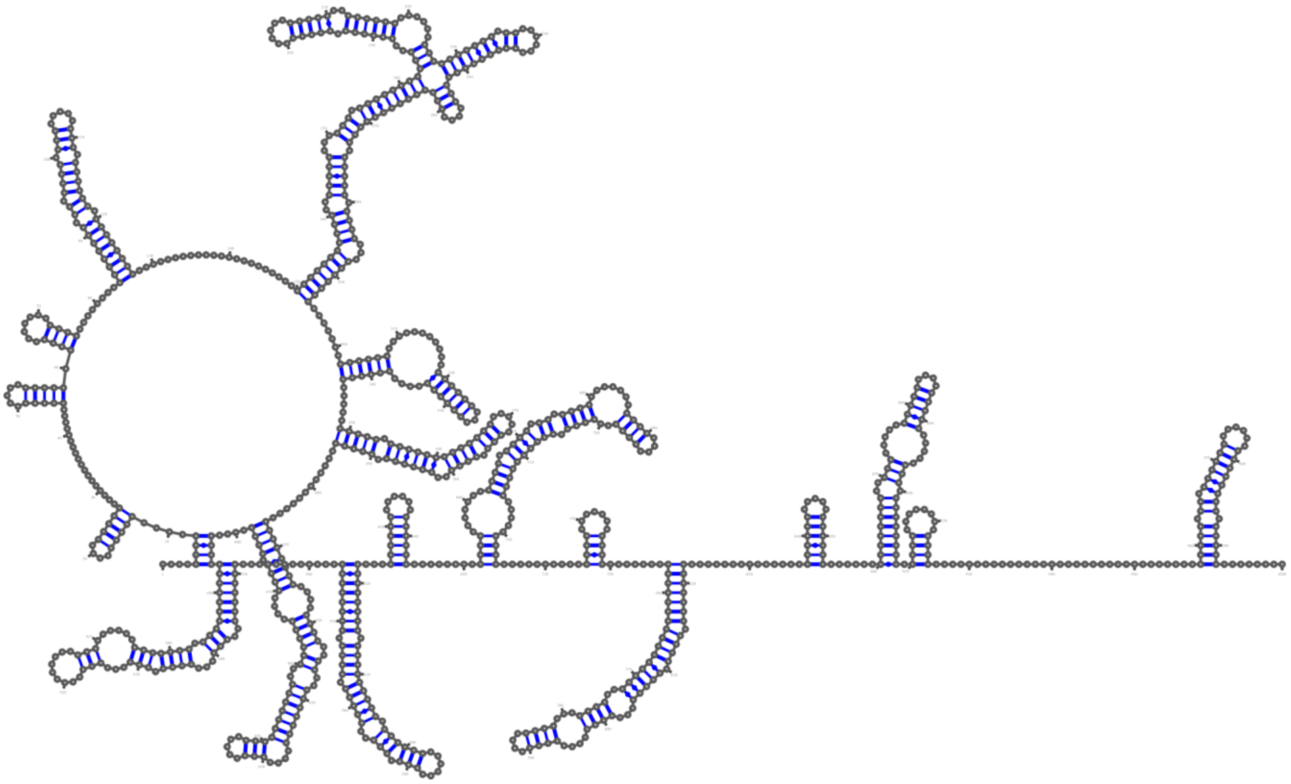

**Figure S2: Secondary Structure of g1-1000.** Structure taken from published SHAPE-MaP data[40] on g1-1000.

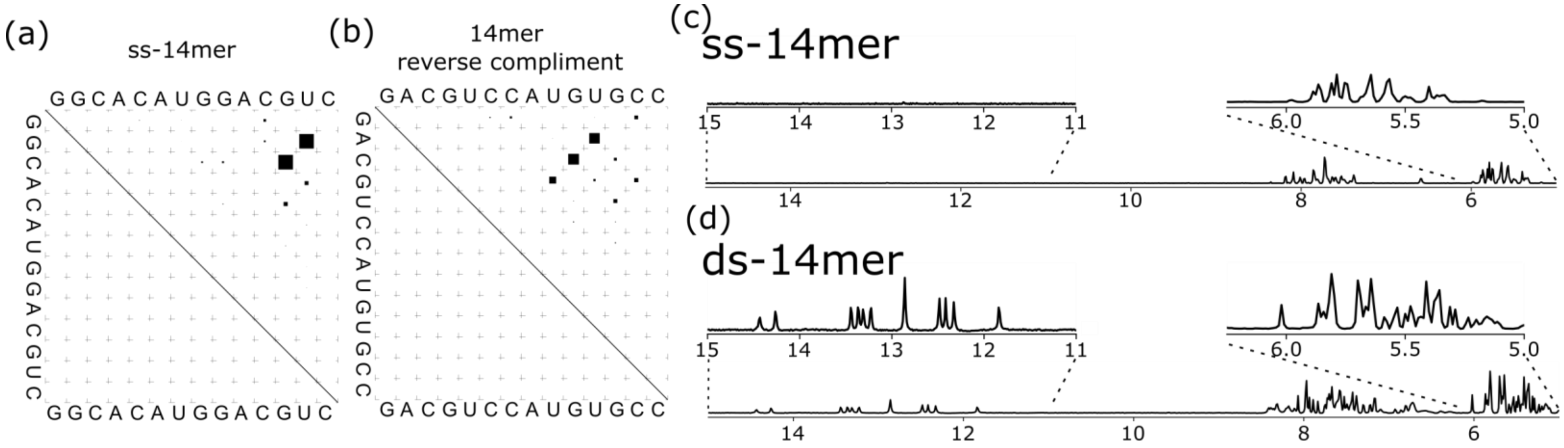


**Figure S3: Characterization of 14mer RNAs.** (a,b) dot plots of the ss-14mer (a) and 14mer reverse complement (b). The region above the diagonal in the dot plot shows the relative probability of base pairs within the equilibrium ensemble. The size of the dots in this section are proportional to the probability of a base pair occurring. The region below the diagonal shows base pairs corresponding to the mean free energy structure. A lack of dots below the diagonal indicates that 14mer and reverse complement have no predicted secondary structure. (c,d) 1H NMR spectra of the ss-14mer (c) and ds-14mer (d) RNAs. The ds-14mer exhibits a pattern of peaks in the 5-6 PPM region that is distinct from the ss-14mer, indicating the presence of double-stranded RNA. In the 11-15 PPM region, peaks that are characteristic of paired bases appear in the ds-14mer spectrum, while none are present in the ss-14mer spectrum.

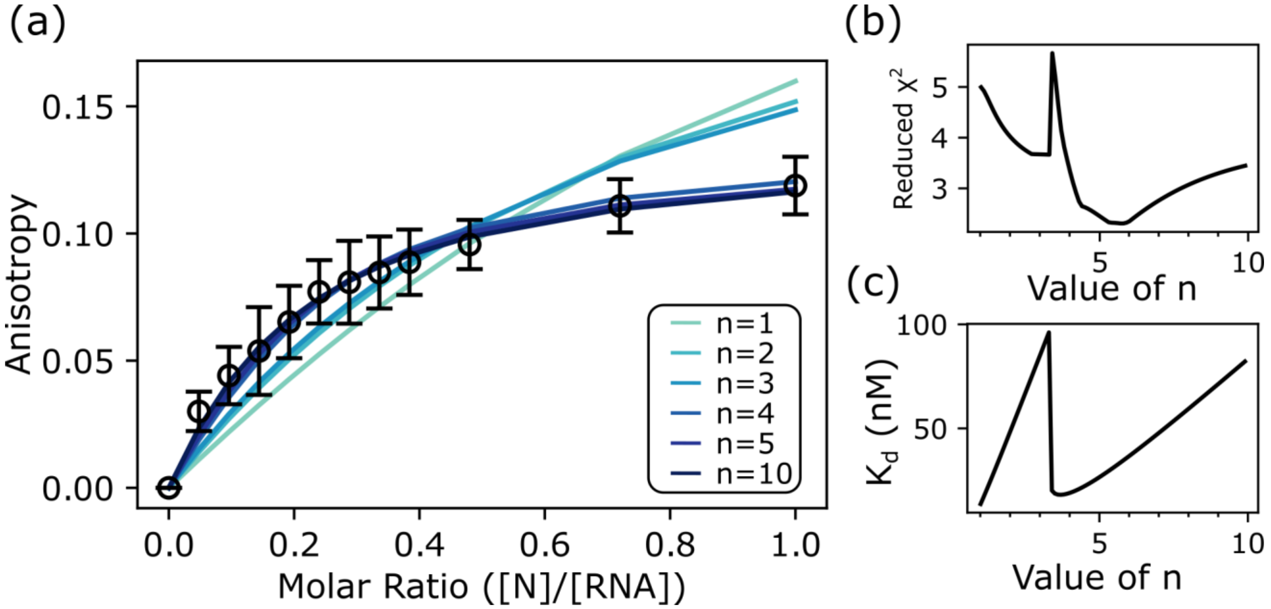


**Figure S4: Analysis of the fluorescence anisotropy curve of FL-N binding to the ss-14mer.** Attempts to fit the curve for this interaction to a 1:1 binding model results in a poor fit (n=1 in (a) above), as N binds multivalently to the RNA[12]. While the underlying mechanism of binding is likely complex, we fit the curve to a simple quadratic model[46] with an additional varied, fixed stoichiometry value (n), representing the number of RNA-binding sites on FL-N. (a) shows example fits at values of n from 1 to 10 (in shades of blue) overlayed with the experimental data (black circles). (b) is an 'error surface' plot of the reduced $\chi^2$ value of the fit as a function of the fixed n value in increments of 0.1, and (c) is a plot of the fit $K_d$ as a function of the fixed n value in increments of 0.1. Using these plots as a diagnostic, we elected to report a fit at n=4 in the main body of the manuscript, following the logic that it is the lowest integer value that fits the data well and reaches a low plateau in the $\chi^2$ error surface. We expect N to have at least 3 RNA-binding sites based on our work here and in previous publications[12], therefore a stoichiometry of 4 is near our expectations, especially when considering additional possible binding sites in N's intrinsically disordered regions.

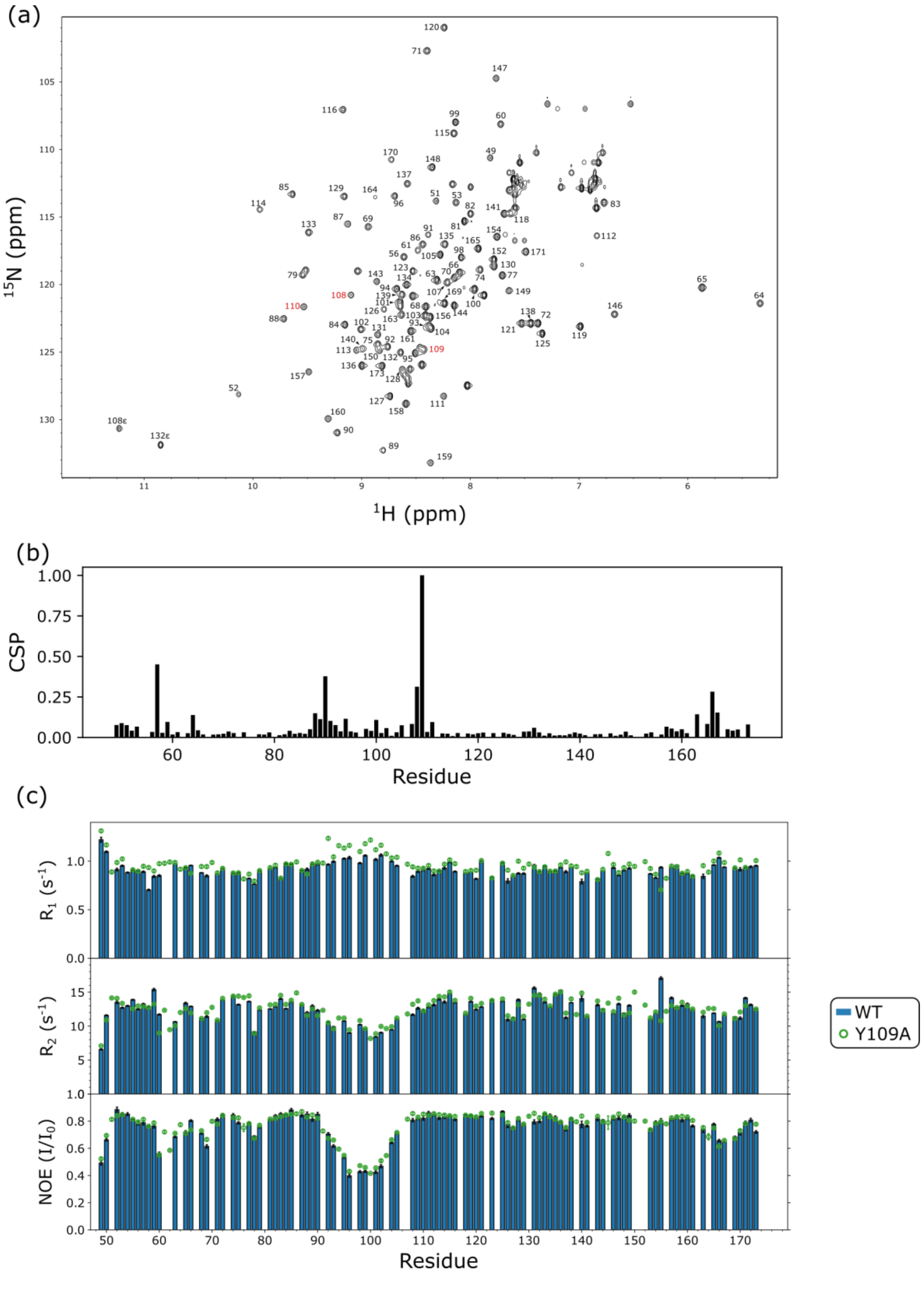


**Figure S5: NMR assignments and dynamics of Y109A NTD.** (a) $^{1}$H-$^{15}$N HSQC spectrum collected at 25°C and a field strength of 800 MHz of the Y109A NTD labeled with assignments. A109 and the resides adjacent to it in sequence are indicated in red. (b) chemical shift perturbations induced by the Y109A mutation relative to the WT NTD. (c) $^{15}$N $R_1$ (top), $^{15}$N $R_2$ (middle), and {$^{1}$H}-$^{15}$N Heteronuclear NOE (bottom) measurements for the NTD collected at a field strength of 800 MHz. Values for the WT NTD are shown as blue bars, and the Y109A NTD as green circles. Comparing the WT to the Y109A NTD reveals their dynamic properties are largely the same, suggesting that the mutation does not have any global impact on the domain's dynamics. The largest difference is in $R_1$ values, which are larger in the Y109A mutant in the 90-110 region that constitutes the NTD's beta hairpin. This is consistent with a slight increase in flexibility of this region, indicating the Y->A mutation may decrease the rigidity of the hairpin.

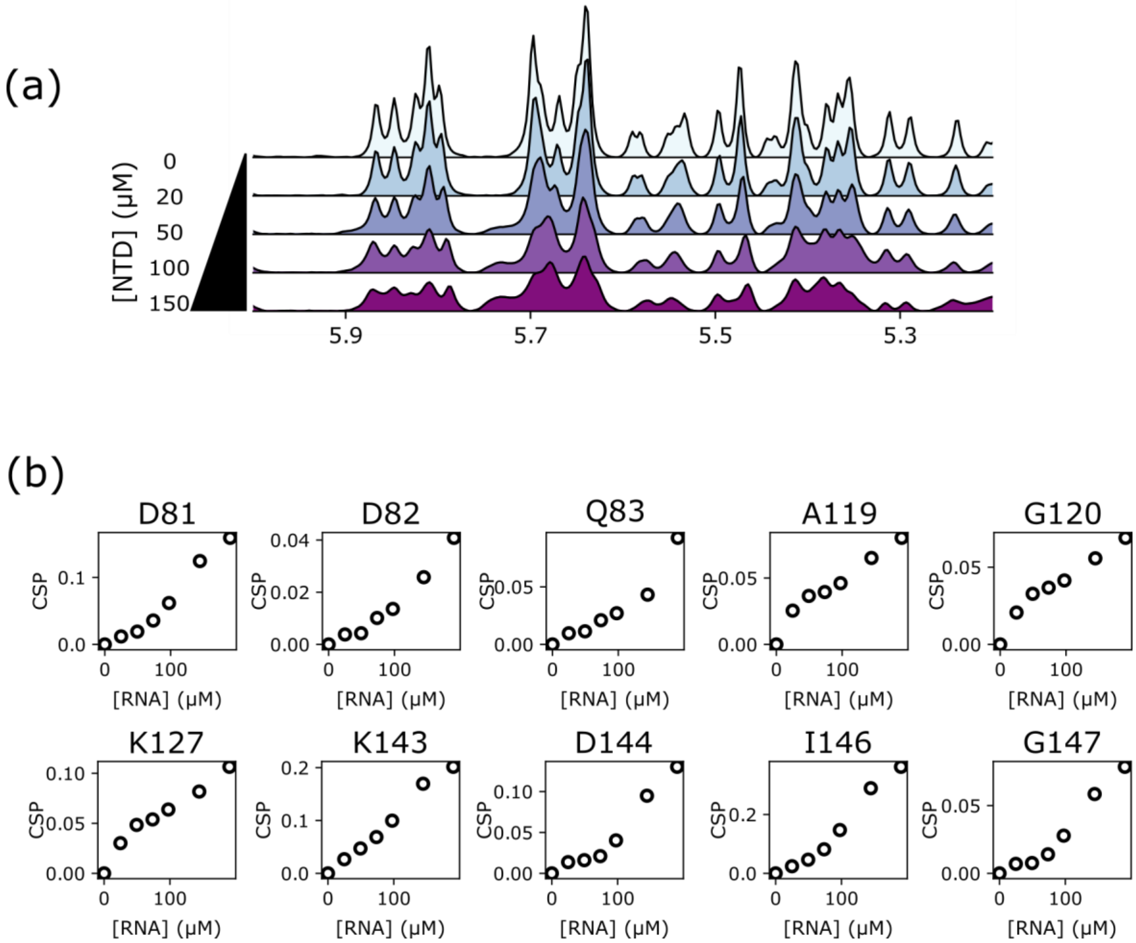


**Figure S6: Additional NMR experiments on the ds-14mer binding to NTD.** (a) $^{1}H$ NMR spectra of the ds-14mer (100μM) in the 5.2-6 ppm region, titrated with the NTD (up to 150μM). Titrations are from the same experiment as Fig. 3h). (b) Chemical shift perturbations for all secondary-site residues from titration of ds-14mer into the NTD**.** Plots are the profiles of CSPs as a function of RNA concentrations for titration of the ds-14mer into 100 μM WT NTD. Residues were selected from the protein based on the linear relationship between CSP and concentration of RNA. These residues cluster together on the NTD structure (See Fig. 4i), indicating that they represent a secondary weak RNA-binding site on the NTD.

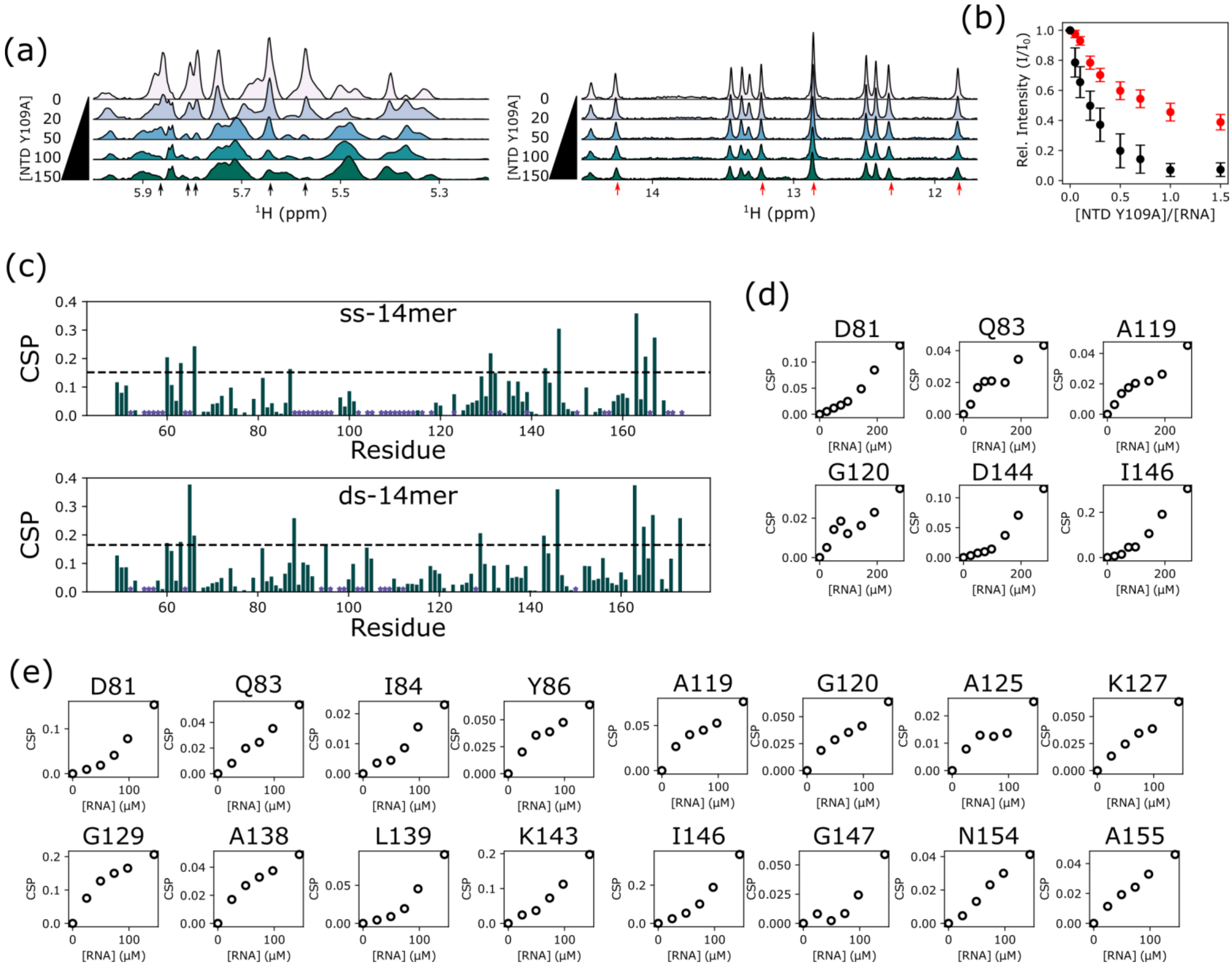


**Figure S7: NMR analyses of Y109A NTD interaction with RNA.** (a) Titrations of Y109A NTD into the ss-14mer (left) and ds-14mer (right). Experimental conditions are the same as in Fig. 3g-i: 100 μM ss- or ds-14mer in a buffer of 20 mM $NaPO_4$ pH 6.5, 20 mM NaCl was titrated with the Y109A NTD, up to a 1.5:1 NTD:RNA ratio. (b) Average peak intensities for the 5 selected peaks (marked with black and red arrows in a), for the titration of NTD into the ss-14mer (black) and ds-14mer (red).(c) CSPs as shown in Fig. 5a,b of the Y109A NTD by residue following titration with the ss-14mer (top) and ds-14mer (bottom). Two standard deviations of the CSP population is drawn as a black dashed line, and residues that disappear due to intermediate exchange are labeled with a purple star.(d) Chemical shift perturbations for residues with linear chemical shift perturbation (as in Fig. 5d) profiles for titration of the ss-14mer into 100 μM Y109A NTD. (e) Chemical shift perturbations for residues with linear chemical shift perturbation (as in Fig. 5f) profiles for titration of the ds-14mer into 100 μM Y109A NTD.

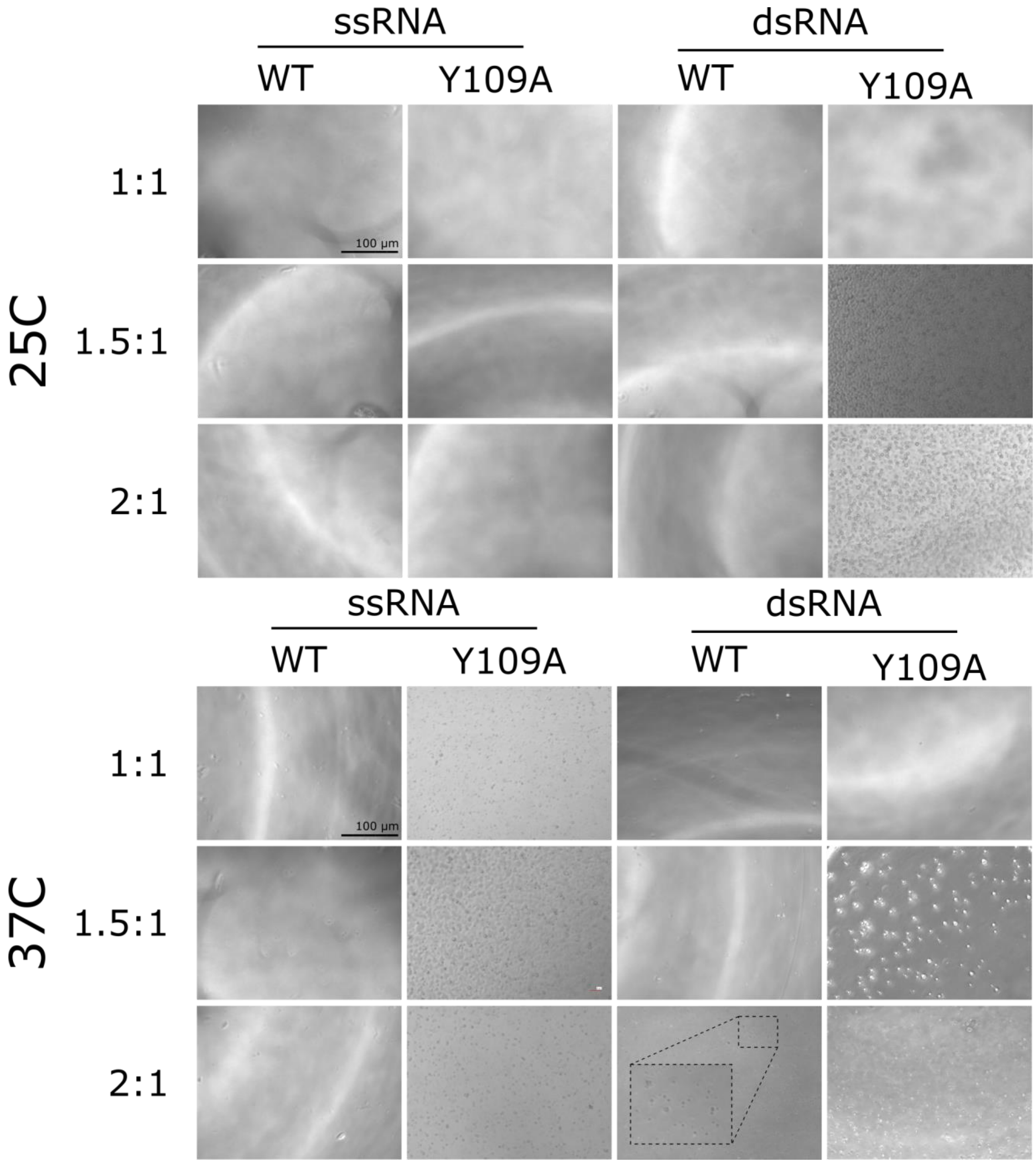


**Figure S8: Phase separation of the NTD at NMR conditions.** Brightfield images of WT and Y109A NTD at NMR conditions (100 µM NTD, 20 mM $NaPO_4$ pH 6.5, 20 mM NaCl), with 100, 150, and 200 µM of RNA (ss- and ds-14mer) added. Images in the top panel are collected at 25°C (used for our NMR experiments) and those in the bottom panel are at 37°C. Scale bar shown in the top left of both panels is 100 µm, and scale is the same for all images.
